# Effects of somatic mutations on cellular differentiation in iPSC models of neurodevelopment

**DOI:** 10.1101/2022.03.04.482992

**Authors:** Pau Puigdevall, Julie Jerber, Petr Danecek, Sergi Castellano, Helena Kilpinen

## Abstract

The use of induced pluripotent stem cells (iPSC) as models for development and human disease has enabled the study of otherwise inaccessible tissues. A remaining challenge in developing reliable models is our limited understanding of the factors driving irregular *in vitro* differentiation of iPSCs, particularly the impact of acquired somatic mutations. We leveraged data from a pooled dopaminergic neuron differentiation experiment of 238 iPSC lines profiled with single-cell and whole-exome sequencing to study how somatic mutations affect differentiation outcomes. Differentiation was tracked at three time points corresponding to neural progenitors, early neurons and mature neurons. We found that deleterious somatic mutations in key developmental genes, notably the *BCOR* gene, are strongly associated with failure in dopaminergic neuron differentiation, with lines carrying such mutations also showing larger proliferation rate in culture. We further identified broad differences in cell type composition between failed and successfully differentiating lines, as well as significant changes in gene expression contributing to the inhibition of neurogenesis, a functional process also targeted by deleterious mutations in failed lines. Our work highlights the need to routinely measure the burden of deleterious mutations in iPSC lines and calls for caution in interpreting differentiation-related phenotypes in disease-modelling experiments.

## Introduction

Induced pluripotent stem cells (iPSC) are widely used to model human diseases as they can differentiate to other cell types, including those from tissues that are otherwise not accessible. However, *in vitro* differentiation is subject to substantial technical and biological confounders that often lead to variable differentiation outcomes, a challenge to scaling and interpreting results from these models of disease ^1^. The underlying reasons for this variability are not well understood, but different factors have been proposed: protocol optimisation ^2^, culture maintenance ^3^, passage number ^4^, molecular determinants ^5^, inter-laboratory variation ^6^, cell line intrinsic properties ^7^ or the loss of iPSC heterogeneity in culture ^8^.

The genetic background of an individual has been shown to account for 8-23% of phenotypic variation in iPSCs ^7^. While this donor effect was driven primarily by common variants, rare variants and somatic mutations acquired either *in vivo* in the parental tissue or *in vitro* during the iPSC reprogramming process and culture maintenance ^9^ are likely to also contribute to the observed variation. For example, it has been shown that sub-clonal cancer-associated mutations might provide a growth advantage to stem cells in culture, given their increase in frequency ^10^. Also, the reprogramming of parental cells, such as skin-derived fibroblasts, can act as a bottleneck, leading to variants increasing or decreasing in frequency in the resulting population of iPSCs ^11^. This can be particularly pronounced if the parental cells contain a higher-than-average amount of mutations, as can be the case with skin-derived, UV-exposed cells. Although such acquired mutations might not cause a phenotype in iPS cells, they have the potential to impact specific differentiated cell types ^12^, altering both their functionality and the overall cell type composition. Still, the contribution of somatic mutations to cellular differentiation has not been systematically explored.

In this study, we hypothesised that somatic mutations in individual iPSC lines might affect their ability to differentiate. To test this hypothesis, we leveraged data from the HipSci project ^7^ and analysed differentiation outcomes from four independent experiments producing different iPSC-derived cell types: dopaminergic neurons, DN ^8^, macrophages ^13^, sensory neurons ^14^ and definitive endoderm tissue^15^ (**Fig. 1A**). We analysed the exome-wide burden of mutations acquired *in vitro* during reprogramming and more generally, all deleterious variants in the iPSCs, which also include germline variants and mutations acquired *in vivo* (**Fig. 1B**). Leveraging single-cell transcriptomics from the dopaminergic neurons ^8^, we studied how deleterious mutations influence the proliferation rate of iPSC lines during differentiation, shifting cell type composition and gene expression that compromised the success of differentiation.

**Figure 1.**
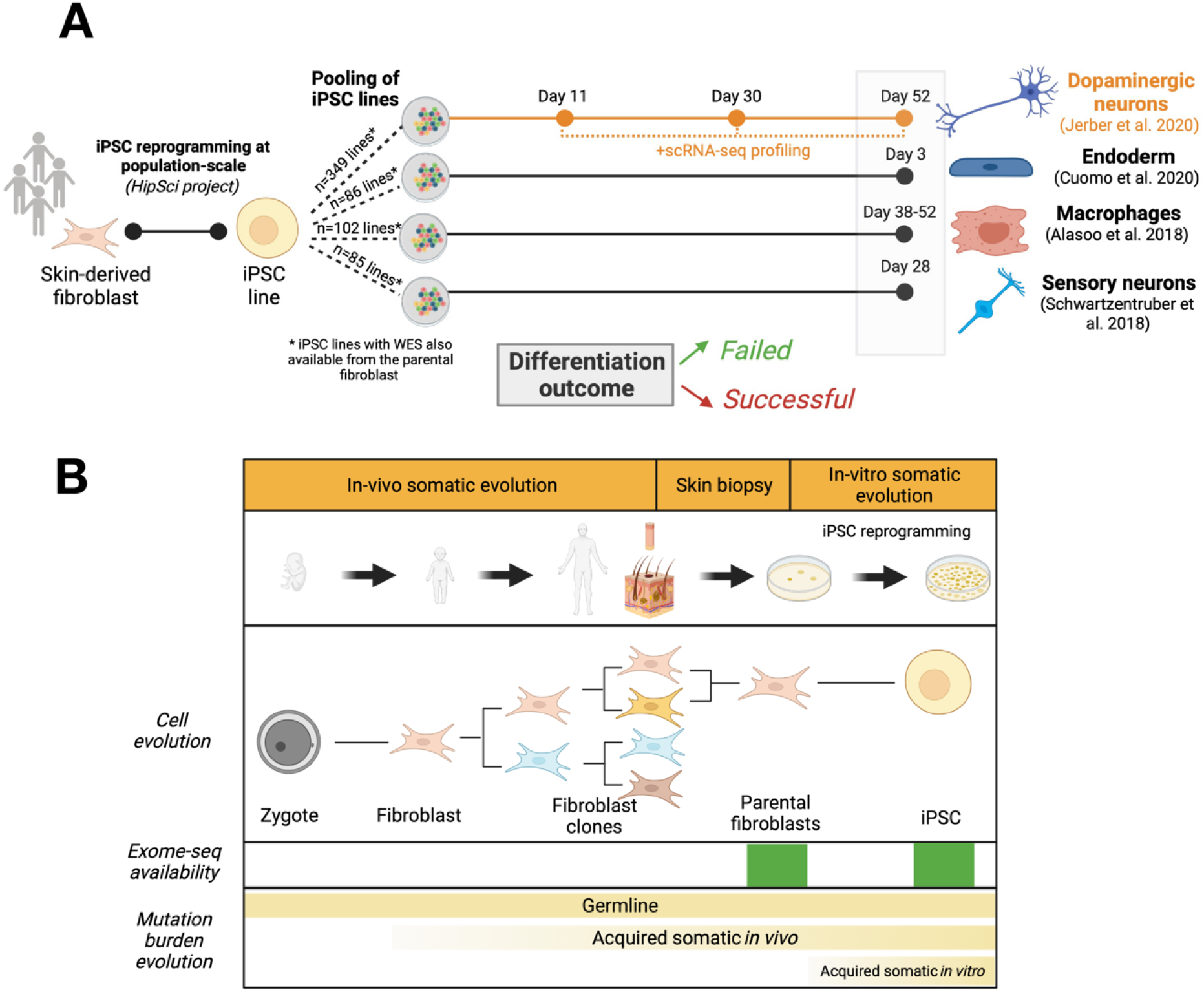
Studying the cellular basis of iPSC differentiation. **a**, Experimental design of differentiations: Fibroblasts derived from the skin of healthy patients were cultured and reprogrammed into iPSC lines (HipSci Project). We leveraged four existing iPSC differentiations that used HipSci lines to derive dopaminergic neurons (N=128, nPred=349), macrophages (N=102), sensory neurons (N=85) and endoderm (N=86). In the former three, lines were classified as having a failed or successful outcome upon the fraction of target cell type observed at the end of the differentiation. As for the endoderm, the differentiation efficiency was given as an average pseudotime at day 3, ranging from 0 (pluripotent state) to 1 (differentiated state) (Cuomo et al. 2020). From the dopaminergic dataset, we used the observed and predicted outcome per line from (Jerber et al. 2020). Cells from this differentiation were profiled at single-cell level at three time-points (day 11, day 30, day 52) aiming to observe progenitors, young neurons and mature neurons. **b**, Characterisation of the iPSC mutational burden. The variants carried by iPSC lines can be classified as germline or somatic mutations upon their origin. Those variants acquired during fibroblast clonal evolution, previous to the skin biopsy, are referred as acquired mutations *in vivo*, while those variants acquired during or after iPSC reprogramming are referred as acquired mutations *in vitro*. We leveraged whole-exome sequencing data from 832 iPSC cell lines to evaluate the genetic determinants on several iPSC differentiations, where germline and somatic mutations were a priori indistinguishable. Yet, for 384 of the 832 lines, a joint variant calling between the available parental fibroblast and the corresponding iPSC (Rouhani et al. 2021) was performed to detect the somatic mutations that were positively selected during reprogramming as a proxy to identify acquired somatic mutations *in vitro*.

We found that the acquisition of deleterious mutations in developmental genes can compromise the success of iPSC differentiation. Lines with such mutations, notably in the *BCOR* gene, are impaired in their capacity to produce dopaminergic neurons and show a faster proliferation rate in culture. This leads to large differences in cell type composition between failed and successful lines, and changes in gene expression that modulate neurodevelopment. Further, we observed that the proliferation rate of lines is predictive of their cell type composition throughout neuronal differentiation. Our work highlights somatic mutations as an important source of variation in iPSC-based models, and calls for caution when interpreting differentiation-related phenotypes to understand disease.

## Results

### 1. The genome-wide burden of acquired mutations does not explain differentiation outcome

To test whether the overall burden of somatic mutations acquired by iPSC lines *in vitro* influenced their differentiation ability, we studied 384 cell lines (251 individual donors) from the HipSci project. Exomes were sequenced for both the parental fibroblast of donors and their derived iPSC lines ^7^, allowing us to distinguish between variants present already in the donors from those acquired, or positively selected for, in the subsequent reprogramming process (hereon ‘*in vitro*-acquired mutations’) ^11^ (**Methods**) A median of 35 mutations (37 when including CNV) was observed per exome in the iPSCs, of which 14 were annotated as deleterious. In line with previous publications ^16^, half (50.5%) of the mutations (SNVs and dinucleotides) were predicted to be missense or LoF. We then considered differentiation outcomes from four different tissues and cell types derived from the same HipSci donors: dopaminergic neurons (DN; N=128 observed differentiations and N=349 predicted differentiations, see **Methods, Supp. Table 1**) ^8^, sensory neurons (N=85) ^14^, macrophages (N=102) ^13^ and definitive endoderm (N=86) ^15^. For each of the cell types, we split the cell lines to those that differentiated successfully and those that failed, including those with impaired capacity (from hereon ‘successful’ and ‘failed’ differentiators) (**Methods**).

**Table :** Cell type annotation using the gene markers provided by ^8^

**Table: Cell type annotation using the gene markers provided by <sup>8</sup>**
| Num. | Cell type | Abbr. | Gene markers used for cell annotation |
| --- | --- | --- | --- |
| 0 | **Dopaminergic neurons** | DA | *ABCC8, ACOT7, ALDH1A1, AMER3, ARG2, ASB4, BNC2, CADPS2, CALB1, CALB2, CAMK2N1, CCK, CDK14, CDKN1C, CHL1, CHRNA4, CPEB3, CXCR4, DCC, DKK3, DRD2, EBF2, EN1, EN2, EPHA5, ERC2, FGF13, FOXA1, FOXA2, GDAP1, GFRA1, GRIA3, GRIK3, GRP, ICA1L, IGF1, KCNIP4, KCNJ6, KLHL1, KLHL13, LGI1, LMO3, LMX1A, LMX1B, LRRC3B, LRRTM2, LSAMP, LY6H, NETO2, NR4A2, NTSR1, OTX2, PBX1, PRICKLE2, PRKCA, PRL, PRRT4, PITX3, PTPN5, PTPRO, RET, SCG2, SLC10A4, SLC18A1, SLC18A2, SLC18A3, SLC6A3, SNCA, SOX6, TH, TMCC3, TMEFF2, TMEM255A, TUB, VGF, SYT1, SNAP25* |
| 1 | **Astrocytes** | Astro | *S100B, AQP4, GFAP, SLC1A3, SOX9* |
| 2 | **Floor-plate progenitors type 1** | FPP-1 | *ZEB2, DMBX1, HMGA1, HMGB2, LMX1A, FOXA2* |
| 3 | **Serotonergic-like neurons** | Sert-like | *CHGB, FEV, GATA2, GATA3, GCH1, GCHFR, HTR1A, HTR1B, MAOA, MAOB, SLC18A2, SLC29A4, SLC6A4, TPH2, SYT1, SNAP25* |
| 4 | **Ependymal-like cells type 1** | Epend-1 | *STOML3, CCDC153, CDHR4, FOXJ1, DNAH11, TTR, MLF1* |
| 5 | **Proliferative floor-plate progenitors type 1** | proFPP-1 | *MKI67, TOP2A, KIAA1524, ZEB2, DMBX1, HMGA1, HMGB2* |
| 6 | **Floor-plate progenitors 2** | FPP-2 | *ZEB2, DMBX1, HMGA1, HMGB2, LMX1A, FOXA2* |
| 7 | **Proliferative floor-plate progenitors type 2** | proFPP-2 | *MKI67, TOP2A, KIAA1524, ZEB2, DMBX1, HMGA1, HMGB2* |
| 8 | **Unknown cell type 1 (Glial cells)** | Unk-1 | *TNC, SOX2, CDH2, HES1, DKK3, EOMES, RELN, LHX1, LHX5, TP73, CALB2, EBF3, SAMD3, SOX2*<br>(Maybe Cajal-Retzius / hem cortical) |
| 9 | **Floor-plate progenitors type 3** | FPP-3 | *ZEB2, DMBX1, HMGA1, HMGB2, LMX1A, FOXA2* |
| 10 | **Proliferative serotonergic-like neurons** | Pro.Sert-like | *MKI67, TOP2A, KIAA1524, CHGB, FEV, GATA2, GATA3, GCH1, GCHFR, HTR1A, HTR1B, MAOA, MAOB, SLC18A2, SLC29A4, SLC6A4, TPH2, SYT1, SNAP25* |
| 11 | **Unknown cell type 2** | Unk-2 | ? |
| 0,3 | **Neuroblasts (DA-lineage/ Sert-like lineage)** | DA / Sert-like (Day 11) | *NEUROG1, NEUROG2, NEUROD1, NEUROD2, NHLH1, SIM1* |

We then tested for association between the genome-wide mutational burden and differentiation outcomes (**Fig. 2A, Supp. Fig. 1A**), and found no difference between lines that failed and lines that differentiated successfully, neither considering all mutations, deleterious mutations (**Fig. 2A**) or other variant classes (**Supp. Fig. 1A,B**). Similarly, no association was found between mutational burden and endoderm differentiation efficiency, which was defined as a continuous trait (**Fig. 2B**).

**Figure 2.**
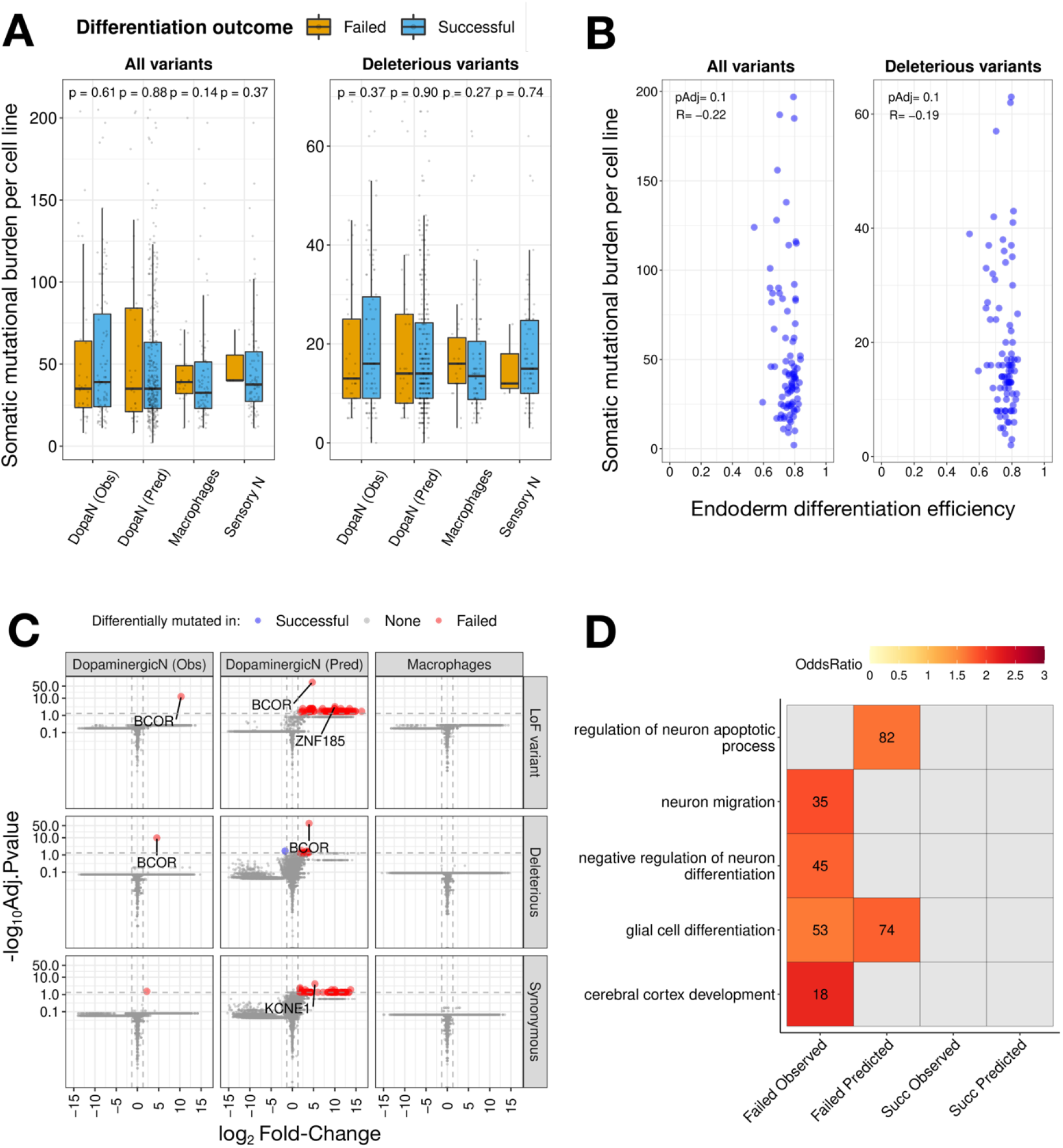
Effect of the mutational burden on iPSC differentiations. **a**, Total (left) somatic mutational burden per cell line - including single nucleotide variants (SNVs), dinucleotides and indels - was not significantly associated with the differentiation outcome observed in macrophages, sensory neurons and dopaminergic neurons (Wilcoxon rank sum test, p>0.05). No association was observed either when considering only deleterious mutations (right). **b**, Similarly, no association (linear correlation, p<0.05) was observed between the total and deleterious mutational burden with the differentiation efficiency in the endoderm (Pearson correlation, pAdj<0.05). **c**, Volcano plot of the differentially mutated genes between the failed and successful lines in dopaminergic neurons (observed and predicted) and for macrophages, tested across loss-of-function (LoF), deleterious (LoF+missense pathogenic) and synonymous variant categories. In red, genes significantly more mutated in failed lines (FC>2.5, Wilcox.test pAdj<0.05) and in blue, genes significantly more mutated in successful lines (FC<1/2.5, Wilcox.test pAdj<0.05). Those genes with the strongest significance are displayed (pAdj<0.001). Only the *BCOR* gene was significantly more mutated in failed lines than successful lines across the variant categories linked to gene damage. **d**, Gene ontology enrichment analysis unravelled the impact of deleterious mutations on neuro-related biological processes (pAdj<0.05) that contribute to the failed differentiation: the negative regulation of neuron differentiation (GO:0045665, hit 45) in failed observed lines or the regulation of neuron apoptotic process (GO:0043523, hit 82) in failed predicted lines. The colour gradient corresponds to the odds ratio (OR) per GO term between failed and successful lines. The number within each tile indicates the hit position of the significant GO term, ordered by decreasing OR within each analysis. Overall, 4 GO analyses were run considering the observed and predicted DN outcomes and for each case, only the top-5% of genes more differentially mutated per outcome were considered (N=933). Neuro-related GO terms that were not enriched in a particular GO analysis are shown in grey.

### 2. The burden of deleterious variants in *BCOR* is linked to differentiation failure in dopaminergic neurons

Mutations that impair the function of developmental genes might have a critical role in defining cell line differentiation efficiency, even when they do not compromise cell survival in culture. We analysed how burden differences in individual genes were linked to the differentiation outcome in the DN dataset. We focused on the burden of deleterious mutations carried by each iPSC line, which includes mutations acquired *in vitro* and *in vivo*, as well as germline variants (**Methods**). We found that only one gene, *BCOR*, was significantly more mutated in failed lines compared to the successful lines (Wilcox.test, p.Adj<0.05, log2FC>2.5) (**Fig. 2C**). This effect was consistently observed with all deleterious variants as well as LoF variants alone, and with both observed (N=183 cell lines; 48 failed, 135 successful) and predicted (N=793 cell lines; 99 failed, 694 successful) DN differentiation outcomes^8^. Importantly, none of the lines that differentiated successfully in culture carried a LoF mutation in the *BCOR* gene, while 22 out of the 48 failed lines carried at least one in *BCOR* (**Supp. Fig. 1C**). The association between the mutational burden in *BCOR* and DN differentiation failure was also seen with the observed differentiation efficiency (pAdj=1.06·10^−8^), defined as the fraction of dopaminergic and serotonergic neurons at day 52, and with the predicted model scores ^8^ (pAdj=7.22·10^−57^) (**Methods**). Although the mechanism for this association is unknown, the *BCOR* gene (a BCL6 repressor) is a known epigenetic regulator^17^ and oncogenic driver gene^18^, as well as a developmental disorder-causing gene^19^, highlighting its role as a key developmental gene. The gene is under strong mutational constraint, with only two predicted LoF SNVs (pLoF) observed in gnomAD (446 expected; LOEUF mutational constraint score 0.141) ^20^. In addition, we did not find any *BCOR* LoF variants among the parental fibroblasts of the iPSC lines, although they could still be present at very low frequency as subclones. Pathogenic mutations in the *BCOR* gene have been found to be recurrently mutated in blood-derived iPSC lines and positively selected for under iPSC culture conditions^11^. In this regard, we cannot determine whether the BCOR mutations driving the impaired DN differentiation originated *in vivo* or *in vitr*o, but they also expanded during iPSC reprogramming.

### 3. Genes disproportionately impacted by deleterious mutations in failed lines are enriched in key neurodevelopmental functions

A fraction of the lines that were classified as failed (26/48, differentiation efficiency < 0.2) do not carry any deleterious *BCOR* mutation, which likely indicates that other genes also contribute to DN differentiation failure but are not identified in our analysis due to limited sample size. To overcome this, we focused on the biological processes that control cellular differentiation and performed a gene ontology (GO) enrichment analysis on those genes that presented the largest mutational burden differences between failed and successful cell lines (top-5% and bottom-5% in fold-change (FC), corresponding to 933 genes for each outcome, **Methods**). We found an overrepresentation of neurodevelopmental GO terms (adjP<0.05) among the set of genes disproportionately mutated in failed lines consistent with the failed differentiation **(Fig. 2D, Supp. Table 2)**, including the negative regulation of neuron differentiation (GO:0045665, hit 45 in observed failed lines) or the regulation of neuron apoptotic process (GO:0043523, hit 82 in predicted failed lines). The most mutated genes (log_2_FC>1.5) contributing to the enrichment of those biological processes in failed lines are well-known disease associations: *NSUN5* in William Beuren Syndrome ^21^, *FOXG1* in FOXG1-syndrome ^22^, *DRD1* in Autism Disorder ^23^ or *ASCL1* in neuropsychiatric disorders ^24^. In other cases, those genes are key regulators of neuron differentiation like *NRBP2* for neural progenitor cell (NPC) survival ^25^, *DMRTA2* for NPC maintenance ^26^, S100A8 involved in microglial inflammation ^27^ or SOX14, essential for the initiation of neuron differentiation ^28^.

Alternatively, we focused on a subset of iPSC lines that were derived from the same donor (replicate pairs, N=49) but were predicted to have a discordant differentiation outcome ^8^. We filtered out those mutations that were shared between the replicates and tested whether the remaining mutations were enriched in two curated gene sets, both including the *BCOR* gene, known to influence organismal development and cellular proliferation: the developmental disorders genes (DDD, N=1,938)^29^ and cancer-associated genes (Cosmic-Tier1, N=558)^30^ (**Methods**). We found an enrichment of LoF variants in cancer-associated genes when comparing the most mutated genes in the failed replicates with the most mutated genes in the successful replicates (empirical p=0.033, proportion test). Also, among failed replicates, we found an enrichment of DDD burden for deleterious (p=0.039) and synonymous variants (p=0.015) (**Supp Fig. 1D**).

### 4. Differentiation failure is driven by increased proliferation rate and *BCOR* LoF mutations

One of the concerns in pooled differentiation experiments, where cell lines are grown and differentiated together, is the potential acquisition of mutations in genes, such as cancer-associated genes, that confer growth advantage to certain lines in culture and lead to an imbalanced representation of the pooled lines. We leveraged the DN experiment ^8^ to assess the *in vitro* growth dynamics of pooled DN differentiation. We reanalysed 846,841 cells from 238 cell lines that were assayed with single-cell RNA-sequencing at three differentiation time points (days 11, 30 and 52, corresponding to progenitors, young neurons and mature neurons, respectively) (**Methods, Fig.1, Supp. Fig. 2A-J**). The dataset consists of 19 different pooled differentiations, including 7 to 24 cell lines per pool (**Supp. Table 3**).

First, we analysed the reproducibility of cell line abundance within each time point across different types of replicates: cell line replicates from different pools or within the same pool, including biological replicates, technical replicates, and 10x channel replicates (**Methods, Supp. Fig. 3A**). Poor correlation was observed between replicates from different pools (R^2^=0.136), which suggests that the specific combination of samples in the pool background can impact the growth dynamics of individual cell lines (**Fig. 3A)**. The correlation improved significantly when comparing biological replicates having the same pool background (R^2^=0.693) and as expected, was highest between technical and channel replicates (R^2^=0.996).

**Figure 3.**
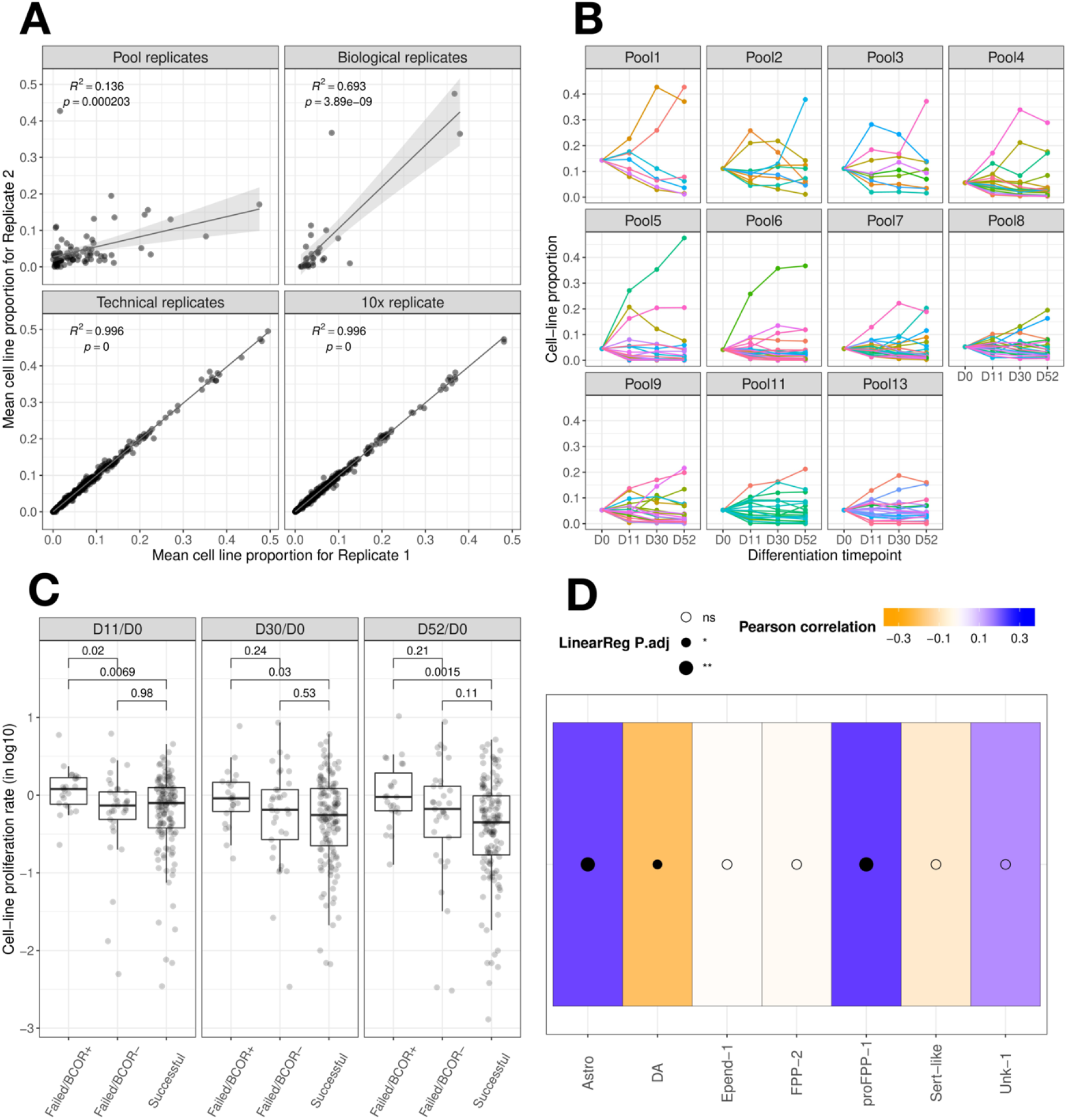
Cell line replicates in pooled differentiation experiments and characterisation of *in vitro* proliferation rates. **a**, Comparison of the mean cell line proportion between replicates of different types: pool replicates, one cell line pooled with different lines; biological replicates, cell lines differentiated with the same pool in independent experiments; technical replicates, cell lines from the same pool differentiated at the same time (different wells of the same plate); 10x replicate, cell lines from the same pool differentiated at the same time (same wells of the same plate). The designation for Replicate 1 and Replicate 2 is random. The p-value and adjusted R-square are obtained from fitting a linear regression. The background of donors in each pool affects the reproducibility of a cell line abundance in culture. Still, the reproducibility was high among independent experiments of the same pool and showed minimal differences between technical and 10x replicates. **b**, Cell line proportion evolution throughout neuron dopaminergic differentiation. Each line corresponds to a cell line in a given pool. Proportions measured at day 11, 30 and 52 are based on cell deconvolution estimates, while at day 0 are based on the equivalent number of cells seeded per line in each pool. We observed a consistent outlier behaviour featured by 1-to-2 lines that proliferate significantly faster during pooled differentiation. **c**, Failed cell lines carrying at least one *BCOR* LoF mutation showed, on average, a higher proliferation rate than neurons that can differentiate successfully in any of the three time points (D11/D0, p=0.0069; D30/D0, p=0.03; D52/D0, p=0.0015, Wilcoxon Rank Sum Test). None of the successful lines carried a LoF mutation in the *BCOR* gene. Each point exhibits the proliferation rate of a cell line in a given pool. **d**, High proliferation rates in cell lines were associated with significant changes on cell type composition at day 52 (linear regression, FDR<0.05), consisting of a depletion of dopaminergic (DA) neurons and an excess of astrocytes and proliferative floor plate progenitors type 1 (proFPP-1). The increasing size of the black-filled circle indicates different levels of significant association: p<0.05 (*), p<0.01 (**); while the white-filled circles stand for non-significant associations. The colour for each cell type indicates whether there is a positive (blue) or a negative correlation (orange).

We then evaluated how the cell line proportions changed during the DN differentiation, assuming initial equal amounts of each iPSC line (**Fig. 3B**) (**Methods**). Specifically, we calculated an *in silico* proliferation rate for each line in a given pool by contrasting cell line proportions at specific timepoints to day 0. Although the Day52/Day0 proportion remained constant for most of the lines (mean=1.0, 95% confidence interval (CI) 0.8-1.2) (**Supp. Fig. 3B**), almost every pool contained at least one overrepresented line (up to 3-10x depending on the pool), with a small fraction of lines being underrepresented (down to 7-700x). Interestingly, those lines that failed to differentiate into neurons showed on average larger proliferation rates than successful lines, with the difference being most significant at the last time point (p=2.8·10^−3^, Wilcox-test) (**Supp. Fig. 3C**). When we correlated this behaviour with the mutational burden, we found that increased proliferation rates were driven by *BCOR* LoF mutations (**Fig. 3C**). Specifically, failed lines with *BCOR* LoF mutations had a significantly higher proliferation rate than successful lines across the time points, a difference that was not observed with failed lines with no *BCOR* LoF mutations. Consistent with this, we found that the *BCOR* gene had the highest ratio of annotated cancer driver mutations (LoF pathogenic Cosmic-Tier1 mutations, N=5) in failed lines compared to successful ones (N=0) (**Methods**). Taken together, these results suggest that proliferation advantage, caused by recurrent somatic mutations in key developmental genes like *BCOR*, has a negative effect on differentiation efficiency.

### 5. Poor differentiation outcomes manifest as shifts in cell type composition already present at the progenitor stage

Next, we studied how early in the differentiation process cell type composition differences between failed and successful lines appear (**Methods**). To that purpose, we processed 119 10x samples (**Supp. Table 4**) that underwent quality control filtering (**Supp. Fig. 2B-D**), dimensionality reduction, batch correction^31^ and Leiden clustering (**Methods**). In order to better characterise the cell type composition of cell lines across the three time points, we processed and clustered all cells in the DN dataset together (**Methods**), in contrast to the original study ^8^. Using the same literature-curated markers (**Methods, Supp. Fig. 2E-G**), we successfully annotated 10 cell types out of the 12 identified clusters (**Supp. Fig. 2H-J**), compared with the 12 out of the 26 identified in the original study.

We used a negative binomial regression model to evaluate cell type composition changes between failed (N=58) and successful (N=163) lines (**Methods**). The analysis revealed significant shifts in abundance for all major cell types (>2% fraction) as early as day 11, except for floor-plate progenitors type 1 (FPP-1) and ependyma (Epend-1) (**Fig. 4A**). Interestingly, cell lines that failed to generate mature neurons at day 52, showed an earlier commitment to either the dopaminergic (DA, pAdj=9.9·10^−47^) or the serotonergic (Sert-like, pAdj=6,1·10^−14^) fate at day 11, represented by neuroblasts clustering in such cell types. Similar evidence of accelerated neuronal maturation *in vivo*, caused by a heterozygous nonsense mutation in *KMT2D*, has been observed in an iPSC model of Kabuki syndrome ^32^. The overall lower fraction of neurons in failed lines is accompanied by a significantly larger proportion of astrocytes (pAdj=1.5·10^−36^), ependymal-like cells (pAdj=7.7·10^−14^) and of the unknown cell type 1 (pAdj=1.1·10^−19^).

**Figure 4.**
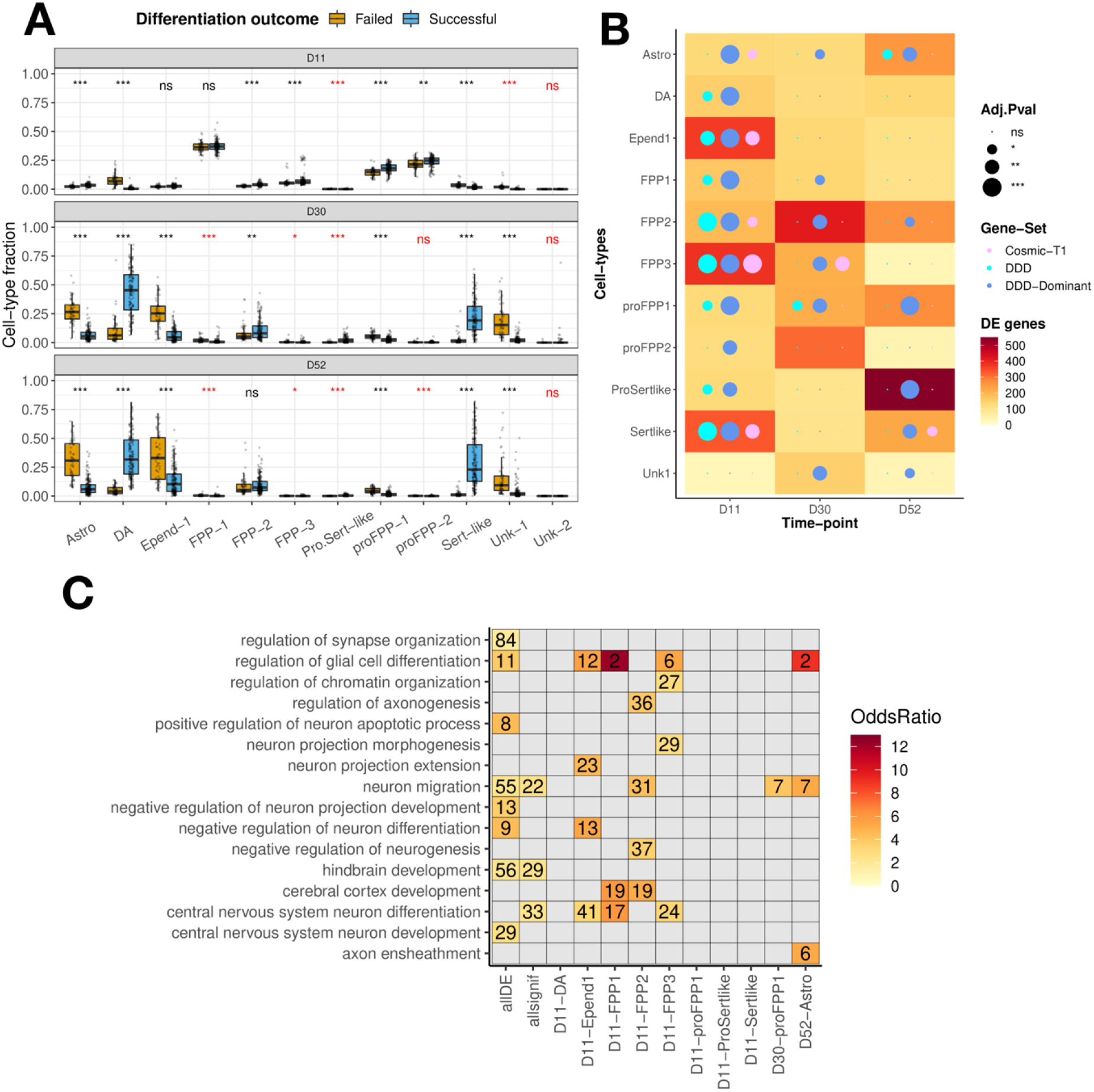
Failed and successful lines showed major differences in cell type composition, gene expression and functional enrichment. **a**, Major cell types showed significant differences in composition as early as in day 11 (negative binomial regression, Methods), except for floor-plate progenitors type 1 (FPP-1) and ependymal-like cells (Epend-1). Neuroblasts from failed lines showed an earlier commitment to dopaminergic (DA) or serotonergic-like (Sert-like) fate, which was notoriously reversed throughout neuron maturation (days 30, days 52). The significance for each test is displayed in red for those cell types with less than 2% of abundance within a given time point. The significance level of each test was indicated as follows: pAdj<0.05 (*), pAdj<0.01 (**), pAdj<0.001 (***), after multiple test correction using the Benjamini & Hochberg method. **b**, The heatmap illustrates the magnitude of differential expression during differentiation between the matched cell types of failed and successful lines. At the progenitor stage (day 11), most of the differentially expressed genes are enriched in developmental disorder genes (DDD, cyan), especially when considering those that act in a dominant fashion (DDD-dominant, blue) (Chi-squared test, Methods). Also, some of the cell types show an enrichment in cancer-associated genes (Cosmic-Tier 1, magenta), particularly strong in the case of floor-plate progenitors type 3. The significance level for the gene set enrichment is indicated by the increasing size of the filled circles: pAdj>0.05 (ns), pAdj<0.05 (*), pAdj<0.01 (**), pAdj<0.001 (***). **c**, Functional enrichment of biological processes (hypergeometric test for GO term association, pAdj<0.05 and FC>1.5) among the differentially expressed (DE) genes. Only the neurodevelopmental and chromatin-related processes are illustrated. Several tests were run, either aggregating all differentially expressed genes (allDE), all the DE genes with an overrepresentation of DDD genes (allsignif) or any of the individual time point and cell type combinations. Several processes involving neuron development and neuron maturation were affected as previously observed in failed lines with differential mutational burden.

In agreement with such observations, iPSC lines with deleterious *BCOR* mutations also presented an altered cell type composition compared to lines without *BCOR* mutations, with a significant depletion of neuronal cell types (DA, p.Adj=5,3·10^−6^; Sert-like, p.Adj=8.9·10^−8^, Wilcoxon rank sum test) accompanied by a significant excess of astrocytes (p.Adj=1.3·10^−5^), proliferative floor-plate progenitors (FPP-1, p.Adj=1.9·10^−3^) and one unknown cell type (Unk-1, p.Adj=6.9·10^−3^) (**Supp. Fig. 3D, Methods**). Similarly, we found an association between the proliferation rate and the abundance of those cell types, with lines proliferating faster showing a depletion of DA neurons (p.Adj=0.01, Pearson correlation) and an excess of astrocytes (p.Adj=0.004) and proliferating progenitors (pro-FPP1, p.Adj=0.004) (**Fig. 3D, Methods**). This suggests that differentiation failure is a consequence of a global, early shift in cell type composition among cell lines that proliferate faster and carry damaging mutations in key developmental genes such as *BCOR*.

### 6. Functional enrichment of gene-expression is consistent with mutational differences between failed and successful lines

We then tested whether differences in cell type composition between differentiation outcomes also manifest in their gene expression. For this, we performed a differential gene expression (DE) analysis between failed and successful cell lines within cell types and time points (**Methods**). We identified between 50 to 500 DE genes per test (**Fig. 4B**) that, importantly, were not correlated with the number of cells observed per outcome (**Supp. Fig. 4A**). While the number of DE genes was relatively constant throughout differentiation in certain lineages (dopaminergic, astrocytes, floor plate progenitors type 1), others showed time-point-specific differences. Remarkably, in most of the cell types at the progenitor stage, the DE genes included a significantly larger proportion of DDD genes than non-DE ones (adj.P<0.05, chi-squared test), in particular if only dominant DDD genes were considered. Likewise, cancer-associated genes (Cosmic-Tier1) were overrepresented among the list of DE genes in five cell types at day 11, one at day 30, and another at day 52 (**Fig. 4B**).

We then performed a GO enrichment analysis (**Methods**) on the ten cell types with a significant proportion of differentially expressed DDD genes. We found several biological processes related to neurodevelopment among the top-25 enriched terms (adj.P<0.05, ordered by odds ratio) (**Fig. 4C, Supp. Table 5**), including the regulation of glial cell differentiation (GO:0045685, day 11) and cerebral cortex development in progenitors (GO:0021987, day 11), as well as the neuron projection extension in ependymal-like cells (GO:1990138, day 11). When aggregating all the detected DE genes in the analysis (any cell type), we found processes strongly linked to failed differentiation, the positive regulation of neuron apoptotic process (GO:0043525) and the negative regulation of neuron differentiation (GO:0045665). We also analysed the changes in pathway regulation on the seven cell types with an excess of cancer-associated DE genes (**Methods, Supp. Fig. 4B**), and consistently identified hallmarks of proliferation: upregulation of the tumour suppressor P53, activation of MYC targets and exacerbated oxidative phosphorylation. This functional enrichment of DE genes is consistent with that of somatic mutations in failed lines.

### 7. Proliferation rate predicts cell type outlier status of cell lines

Cellular differentiation is a dynamic process with global changes in the composition of cell populations over time. Although differentiation success is usually defined by the final yield of the desired cell type, this can give an incomplete picture of the variability in the differentiation process. In an attempt to characterise such variability, we aimed to identify cell lines that were differentiation outliers in terms of their cell type composition. We analysed all pooled cell lines with confident cell fraction estimates (day 11, N=195; day 30, N=212; day 52, N=246) to evaluate their outlier status. For this purpose, we computed the Z-score per line (**Methods**) for each cell type and time point combination and assigned as outliers those cell lines with |Z-score|>2 (**Fig. 5A**). Under this classification, we identified 175 cell lines that were an outlier in at least one of the combinations (day 11, N=58; day 30, N=82; day 52, N=114), most of which had abnormally large cell type fractions. Only on day 11, 16 cell lines showed abnormally low fractions of progenitors and astrocytes, which were compensated by abnormally large fractions of other cell types. We also observed an overall increase in cell type fraction variability at later stages of differentiation.

**Figure 5.**
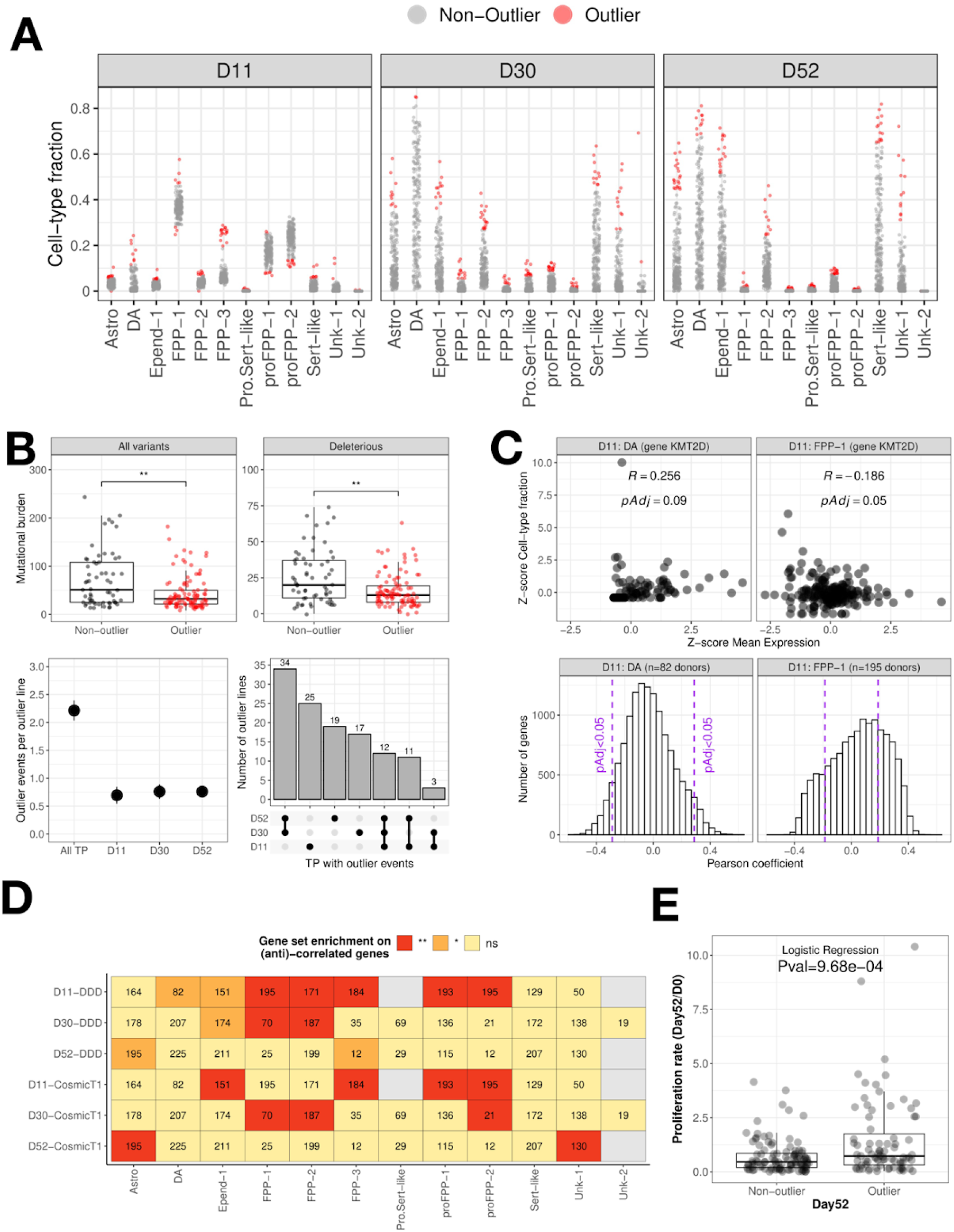
Cell lines in the pooled differentiation displayed common outlier behaviour in cell type composition. **a**, Identification of cell line outliers of cell type composition per time point (dots in red, |Z-score|>2). Most of the outlier cell lines showed an excess of a particular cell type throughout the differentiation, except for some lines with a reduced population of astrocytes and progenitors at day 11. **b**, *upper*: Lines with at least one outlier cell type abundance throughout the differentiation (outlier event) showed a significant reduction in the somatic burden of acquired mutation in vitro than non-outliers (Wilcoxon Rank Sum Test). This is observed considering either all mutations (p=3.6·10^−3^) or just the subset of deleterious ones (p=2.7·10^−3^); *lower*: On average, each outlier line showed 2.21 outlier events per differentiation, but less than 1 event per time point. Also, most of the lines displayed the outlier behaviour either simultaneously at days 30 and 52 (young and mature stage), or just at day 11 (progenitor stage). Each dot represents the mean outlier events while the bars represent the 95% CI. **c**, *upper*: Example illustrating the z-score correlation of the KMT2D gene expression in neuroblasts committing to dopaminergic neurons (DA) with the corresponding cell type proportions at day 11. This analysis was done for all the expressed genes per time point. Each dot corresponds to a cell line; *lower*: The distribution of Pearson correlation coefficients for all genes expressed in the neuroblast populations committing to DA at day 11. We sampled the correlated (and anti-correlated) genes that showed significant associations (pAdj<0.05, purple dashed line) between the expression and the cell type abundance. This distribution was generated for all cell types at each time point. **d**, The abundance of progenitor population was associated with the expression of developmental disorder (DD) and cancer-associated (Cosmic-Tier1) genes. A gene set enrichment (Chi-Squared test, B&H multiple-test correction) was performed between those significantly correlated genes. Results are illustrated in heatmap tiles as follows: not tested due to insufficient cells (grey), pAdj>0.05 (ns, yellow), pAdj<0.05 (*, orange), pAdj<0.01 (**, red). **e**, The mean proliferation rate per line, calculated as the ratio of the cell line proportion between days 52 and 0, was higher in outlier lines than in non-outlier lines (p=9.68·10^−4^, logistic regression).

We further explored the correlation of the somatic mutational burden acquired *in vitro* with the outlier behaviour. Unexpectedly, we observed a significant reduction of burden (p<0.01, Wilcoxon test) in the outlier group (**Fig. 5B**, upper). This difference was observed for both total and deleterious mutations, but when detaching the outlier status per time point, the difference remained significant only at day 30 (**Supp. Fig. 5, Methods**).

To further characterise the outlier behaviour, we calculated the number of times an outlier line shows an abnormal cell type fraction across the differentiation (outlier event, **Methods**). On average, we observed 2.21 outlier events per line (2.03-2.39, 95% CI) with even contributions per time point (**Fig. 5B**, lower). When focusing only on the cell lines profiled at the three time points (N=121 cell lines), we observed that they tend to show outlier events most frequently at the two consecutive latest time points (day 30 and day 52) or just initially (day 11). Only twelve cell lines showed outlier behaviour in all time points.

Although observing abnormal cell type fraction is common among wild-type iPSC lines in pooled experiments, we found that larger proliferation rates at day 52 were associated with the outlier behaviour (p=9.68·10^−4^, logistic regression) (**Fig. 5E**). To identify which genes might be driving this behaviour, we correlated the cell type specific expression with the changes in cell type composition (**Fig. 5C, Methods**). Among the significant associations, including positively and negatively correlated genes, we observed a strong enrichment of DDD genes (p.Adj<0.01, chi-squared test) in most of the cell types at the progenitor stage, at day 11 (**Fig. 5D, Methods**). Similarly, we observed that 7 out of the 9 cell type associations enriched in cancer-associated genes were also enriched among DDD genes, as expected from the significant overlap between the two gene sets (p<2.2·10^−16^, Chi-squared). This suggests that the regulation of developmental genes during early neural induction is critical to determine the progenitor abundance, with further potential to increase the proliferation rate and impact the differentiation success *in vitro*, as shown for the *BCOR* gene.

## Discussion

One of the biggest limitations of iPSC-based disease modelling is our poor understanding, and control, of factors that influence the capacity of cell lines to differentiate successfully and reproducibly. Recently, it was proposed that the large variability associated with differentiation is primarily explained by cell-intrinsic factors^8^, rather than experimental or other technical factors. Genetic variation has previously been shown to drive molecular heterogeneity in iPSCs ^7,33,34,35^, but large-scale studies looking at the role of genetic variation in differentiation are only beginning to emerge. In particular, somatic mutations acquired prior to the differentiation, either *in vivo* during parental tissue clonal evolution or *in vitro* during iPSC reprogramming and culture maintenance, are potential contributors to variation in differentiation efficiency. Here, we present the first attempt to link differentiation outcomes to acquired somatic mutations in human iPSC lines from the HipSci resource, which offered a unique opportunity to study multiple cell types independently derived from the same iPSC lines. Further, with exome sequencing available for both iPSCs and their parental fibroblasts, we were able to focus on the subset of damaging mutations acquired *in vitro*, which are particularly relevant for abnormal differentiation outcomes.

One of the key insights from our work is that although the total burden of acquired mutations in iPSC lines is not predictive of their differentiation outcome, deleterious mutations in the core genes of a given differentiation system can cause unwanted effects on differentiation. This effect is likely not limited to mutations acquired *in vitro*, as mutations and rare variants in the genetic background of the parental cells selected for reprogramming may account for a considerable fraction of differentiation variability, even if not affecting reprogramming directly. In support of this, we found that somatic deleterious mutations in the *BCOR* gene are strongly associated with differentiation failure in human dopaminergic neurons. The effect was seen with 183 observed differentiations as well as 793 predicted differentiation outcomes of iPSC to dopaminergic neurons^8^. The sensitivity to *BCOR* deleterious mutations is supported by the strong selection against predicted LoF variants in the Genome Aggregation Database^36^. Further, a high prevalence of acquired *BCOR* mutations was previously found in blood-derived iPSC lines, and it was shown that they likely arose after reprogramming through positive selection for BCOR dysfunction ^11^. In our dataset, damaging *BCOR* variants were only observed in the iPSC lines, although they could have originated *in vivo* and be present in parental fibroblasts as subclones at very low frequencies that later underwent positive selection *in vitro*.

*BCOR* is a key transcriptional regulator during embryogenesis. It is part of a specific type of polycomb repressive complex that mediates transcriptional repression through epigenetic modifications of histones ^18^ and has been shown to have a key role in regulating the pluripotent state and differentiation (Z. Wang et al. 2018). Like many other chromatin-related genes, *BCOR* is annotated both as a developmental disorder gene and a cancer driver gene ^30,37,38^ we observed that failed lines that carried deleterious *BCOR* mutations showed significantly larger proliferation rates than lines that differentiated successfully, suggesting that monitoring cell line proliferation rates prior to differentiation may be an effective way to screen out lines that will not differentiate correctly ^39^.

Although the *BCOR* gene was not listed in the expression signature of failed differentiation for specific iPSC populations (N=184 lines) (Jerber et al. 2021), we observed that *BCOR* expression in dopaminergic neurons was negatively associated with final neuron abundance, and was also linked to the abundance of progenitor populations earlier in the differentiation. As an epigenetic modulator of stemness and differentiation, it is unclear whether the expressivity of *BCOR* LoF mutations already manifests at iPSC, at precursor stage or at both, potentially redirecting neuron differentiation to astrocytes and ependymal-like cells.

Despite the strong association of *BCOR* mutations with differentiation failure, not all of the failed lines carried damaging variants in that gene, suggesting other genes are involved as well. For this reason, we also analysed the most differentially mutated genes in each differentiation outcome to pinpoint the biological processes that were affected by deleterious mutations. Interestingly, genes that were mutated only in failed lines were enriched in neurodevelopmental processes such as the negative regulation of neuron differentiation or the regulation of the neuron apoptotic process. Similarly, when comparing the top differentially mutated genes per outcome between discordant replicate pairs, failed lines showed an excess of deleterious mutations in DDD genes and an excess of LoF mutations in cancer-associated genes.

To better characterise the cell type composition dynamics throughout the DN differentiation process, we introduced a critical modification to the analysis in the original study ^8^. Specifically, we clustered all cells at once, rather than per time point. While we lose some granularity in the definition of cell types, this approach allowed us to observe cell type composition changes per line across neuron lineages, tracing the commitment of neuroblasts to young and mature neurons. We identified a larger fraction of neuroblasts committing to dopaminergic and serotonergic neurons in failed lines, suggesting an accelerated maturation at the progenitor stage potentially linked to the proliferative phenotype. In that scenario, failed lines could reach faster the number of cell cycles required for differentiation to begin after neural induction, promoting the early production of neuroblasts with a defective neuronal commitment.

Finally, we compared the extent of differential gene expression per cell type across time points and conditions between failed and successful lines. With these comparisons, we sought to identify the key regulator genes across the different stages of neurodevelopment and across different biological processes. Many cell types at the progenitor stage (day 11) showed an enrichment of DE genes corresponding to key developmental genes, either DDD or cancer-associated. In those cell types, the differentially regulated neurodevelopmental processes clearly overlap with the functional processes affected by deleterious mutations in failed lines. We hypothesise that among those DE genes, there is a potential list of new DDD candidates, whose clinical significance should be evaluated. Moving beyond the outcome definition, we found that those genes whose expression is associated with the abundance of progenitor populations at day 11 are also enriched in DDD genes.

Any differentiation process involves a dynamic evolution of cell types, which does not necessarily fit into a failed or successful outcome based on an arbitrary threshold. To avoid overlooking other relevant changes, we analysed outlier behaviour in cell type composition. We found that 64.3% of lines in pooled experiments occasionally display abnormal cell type fractions during the differentiation process. This outlier behaviour reflects the large variability in cell type composition during *in vitro* pooled differentiations, likely resulting from the combination of donor effects, cross-interactions among pooled lines and stochasticity. The periodicity of outlier events suggests that they tend to happen either consecutively in the last two time-points or only at the first one, which can be explained by the experimental design, as cells were only passaged at day 20. Even more importantly, we found that outlier behaviour was strongly associated with larger proliferation rates in cell lines, possibly implying that acquired mutations in other genes that also increase proliferation activity could be behind the abnormal cell type composition. However, since the mutational recurrence in all other genes was substantially lower than in *BCOR*, we were not sufficiently powered to detect population-level evidence for this possibility. Finally, we did not observe a higher burden of acquired mutations in outlier lines when compared to non-outlier lines, but instead the opposite. These observations suggest that while individual deleterious mutations can define differentiation outcomes, the determinants of outlier behaviour during neuronal differentiation are likely more varied.

In summary, our study demonstrates that although iPSC models are an excellent tool for studying neurodevelopment and developmental disorders, results from differentiated cell types should be interpreted with caution. We studied a large number of iPSC lines derived from healthy individuals and observed that deleterious mutations in genes known to cause developmental disorders cause differentiation defects via transcriptional and cell type composition changes during neuronal differentiation. Our work highlights somatic mutations as a significant source of variation in iPSC-based disease models, and further emphasises the importance of comprehensively assaying the genomes of iPSC lines prior to their experimental use.

## Supplementary Figures

**Supp. Fig 1.**
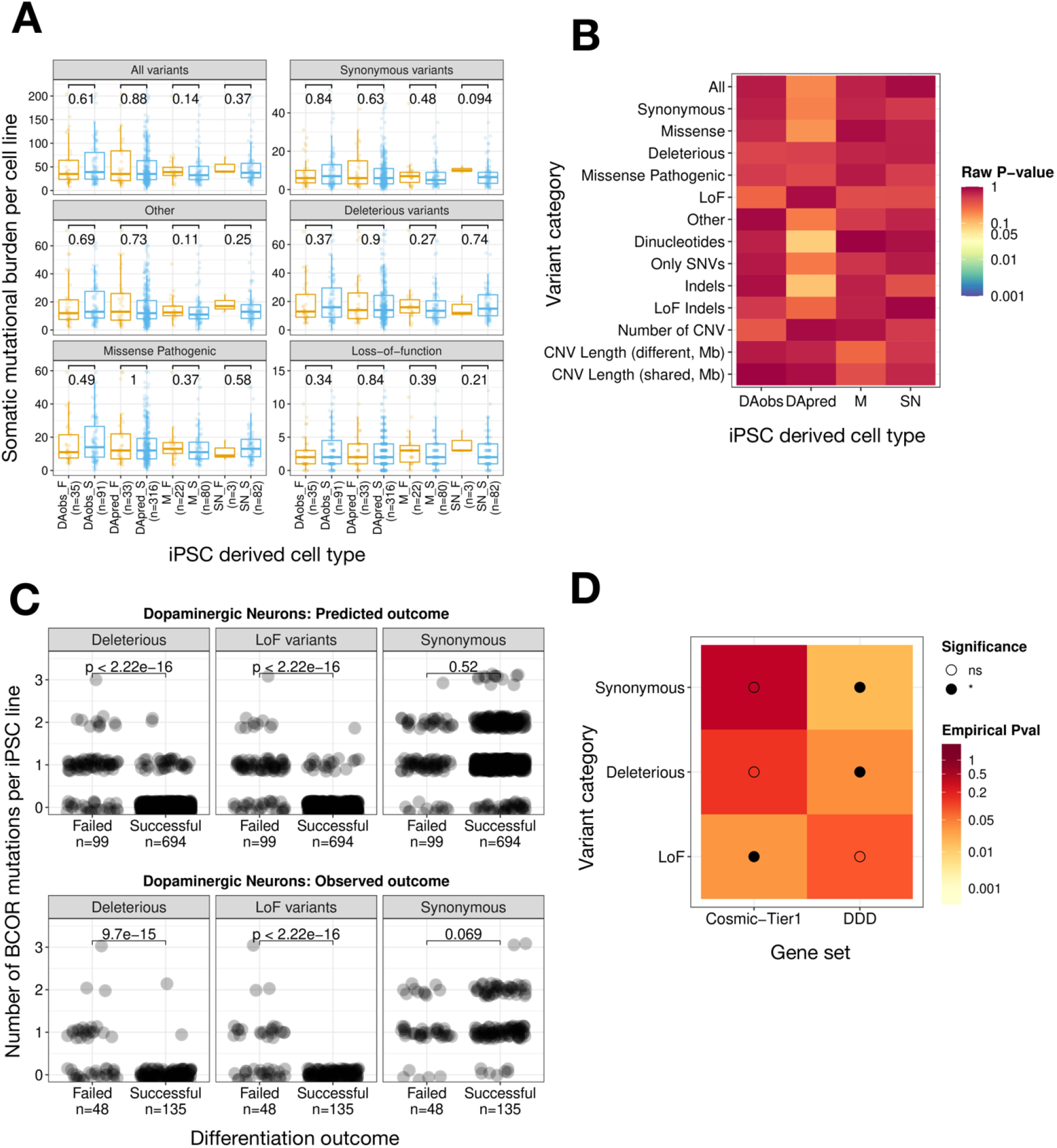
**a**, No significant mean differences on total number of mutations acquired *in vitro* (Wilcox rank sum test, FDR<5%) were observed between the two differentiation outcomes (failed vs successful) of the three different iPSC-derived cell types (dopaminergic neurons - observed and predicted-, macrophages and sensory neurons). We also tested such differences of burden on six variant classes: total variants excluding CNVs; synonymous variants and others (coding, non-coding and unannotated), deleterious variants (union of LoF and missense pathogenic), as well missense pathogenic and loss-of-function variants alone. In none of the 24 tests, the difference reached the statistical significance threshold, even before multiple test-correction. **b**, No association was observed between the binary differentiation outcome (failed or successful groups) and the total number of mutations acquired *in vitro* for any of the tested variant classes. For each mutation category, we fitted a logistic regression and showed the corresponding raw p-values shown in the heatmap. None of the tests reached the statistical significance threshold even before multiple-test correction. We considered all mutations (without CNVs), synonymous, missense (pathogenic and non-pathogenic), deleterious and LoF only, other mutations (coding, non-coding and unannotated), dinucleotides, single nucleotide variants only, indels, indels predicted to be LoF, the number of CNVs, and the region length (in Mb) of shared and different CNVs. **c**, Number of *BCOR* mutations per iPSC line (either LoF, deleterious or synonymous variants) linked to the DN differentiation outcome (observed and predicted). A significant higher burden of damaging variants were observed across failed lines (Wilcox test, p<0.05). None of the successful lines (N=135) in the DN observed outcome carried a BCOR LoF mutation, while 22 out of 48 failed lines have at least one mutation. **d**, Gene set enrichment for developmental disorder (DD) and cancer-associated (Cosmic-Tier 1) genes on the differential mutational burden between discordant replicate lines (N=49 pairs) (Methods). A significant enrichment of LoF burden (nominal p=0.033) was observed in cancer-associated genes, as well as deleterious (nominal p=0.039) and synonymous (nominal p=0.008) burden in DDD genes. The proportion t.test p-value was ranked through 1,000 random gene sets. The colour scale indicates the magnitude of the nominal p-value, while the black-filled circle per tile indicates a significant enrichment (nominal p-value<0.05). The increasing size of the black-filled circle indicates higher significance levels for gene set enrichment: p<0.05 (*), p<0.01 (**), p<0.001(***).

**Supp. Fig 2.**
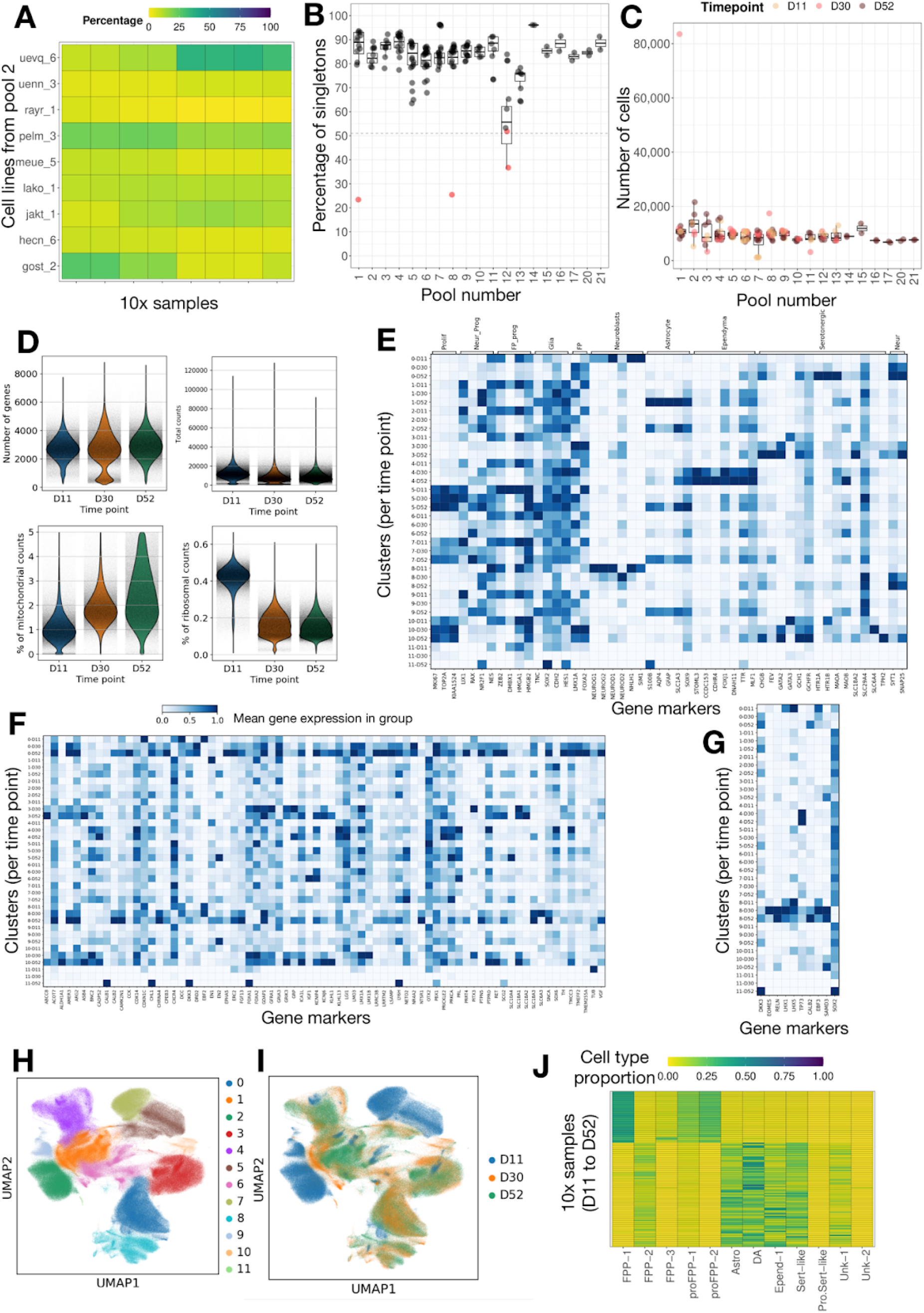
**a**, Cell line proportion for the different 10x samples run for pool 2. Each heatmap row is a cell line and each column corresponds to a 10x sample. In certain samples, one-to-two cell lines consistently account for up to 50% of the cells. **b**, Percentage of singletons confidently assigned to a 10x sample (each dot) of day 52 grouped by pool. As a quality control step, those 10x samples with less than a 50% of singletons were removed from further analysis due to concerns on data integrity. **c**, Number of cell droplets estimated per 10x sample after CellRanger processing. **d**, Quality control metrics after the merging and the cell deconvolution of the 115 10x samples from the dopaminergic differentiation. The median number of genes per cell was below 3,000, while the total count of reads was around 10,000, as expected. Also, cells at day 52 showed a higher average of mitochondrial counts, while at day 11 the higher percentage corresponded to ribosomal counts. **e-g**, Time-specific cell markers expression from ^8^ used to annotate our Leiden clustering. General makers are shown in (e), specific dopaminergic markers in (f) and glial cells in (e). The intensity of blue shows the mean expression of the gene for each cluster-time point group. **h**, UMAP based on gene expression with cells coloured by the annotated cell types. **i**, UMAP on gene expression with cells coloured by sampling time point in differentiation (days 11, 30 and 52). **j**, Cell type proportion in each 10x sample highlights the observed variability on cell type composition throughout differentiation. Each row is a 10x sample ordered by time point, without considering the contribution of individual lines.

**Supp. Fig 3.**
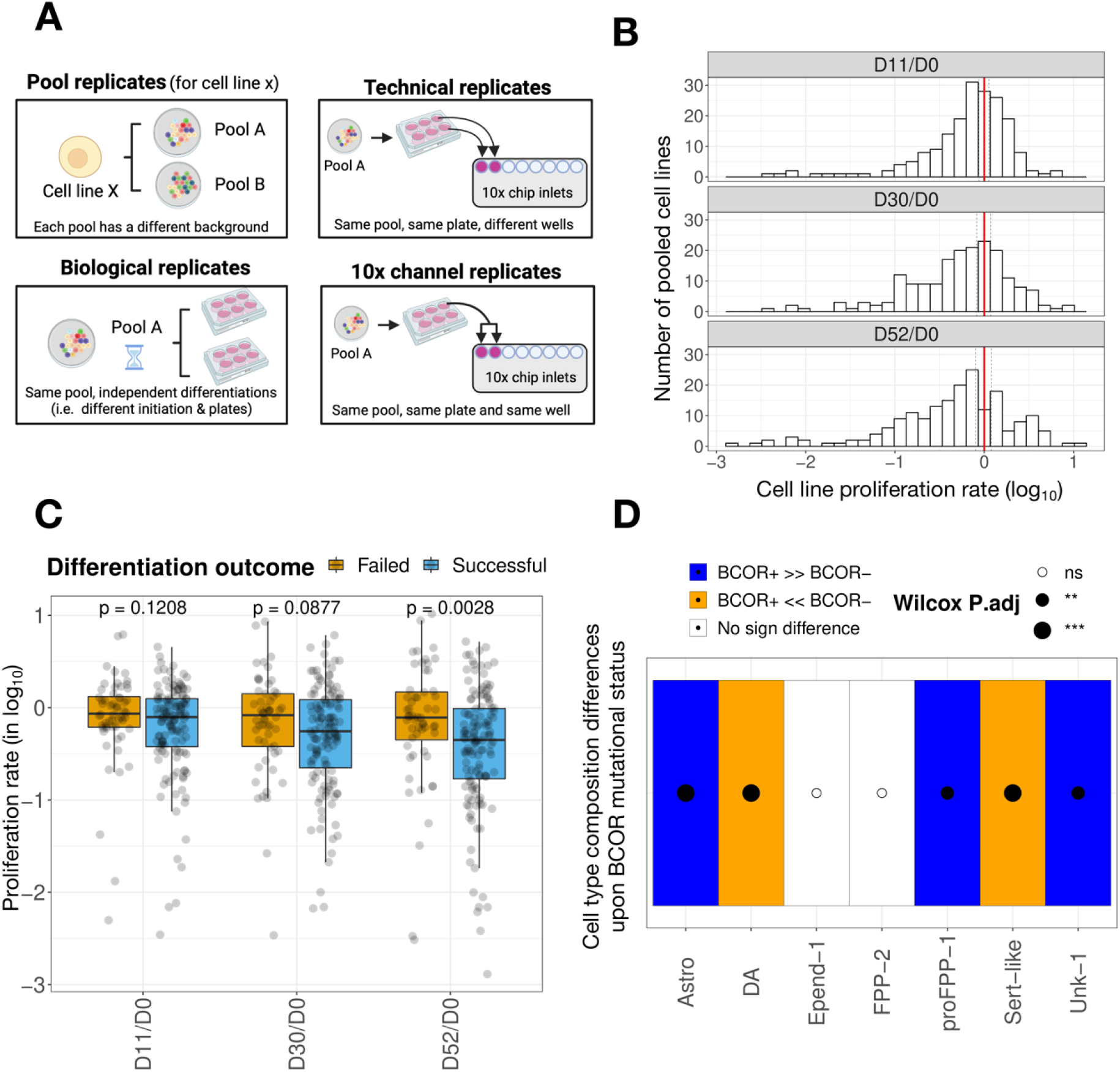
**a**, Illustrative diagram for the four types of replicates considered in DN dataset: pool replicates, referred to cell lines that are included in different pools; biological replicates, referred to lines from the same pool that are differentiated independently (in time and space); technical replicates, referred to lines from the same pool differentiated in the same plate but different wells; and 10x replicates, referred to lines from the same pool, plate and well. **b**, Distribution of the *in vitro* proliferation rate for pooled cell lines (expressed in log_10_) between each differentiation time point and day 0. The vertical red line indicates the mean cell line proportion ratio and the dashed lines the 95% confidence intervals. **c**, Failed cell lines showed on average a larger proliferation rate at day 52 than successful lines (Wilcoxon Rank Sum Test, p=2.8·10^−3^). Proliferation rates are expressed in log_10_. **d**, Cell lines carrying at least one deleterious *BCOR* mutation were associated with cell type composition changes at day 52 with a significant depletion of dopaminergic and serotonergic-like neurons (DA and Sert-like) and an excess of astrocytes (Astro), proliferative floor-plate progenitors type 1 (proFPP-1) and glial cells (Unk-1). The test for association was run for all protein-coding genes (Wilcoxon rank sum test) and corrected for multiple-testing (Benjamini & Hochberg). The increasing size of the black-filled circle indicates higher significance levels for gene set enrichment: p<0.01 (**), p<0.001 (***); while the white-filled circles indicate non-significant association. Cell types with a significantly higher proportion in mutated lines compared to the unmutated ones are labelled in blue. In the opposite scenario, they are labelled in orange.

**Supp. Fig 4.**
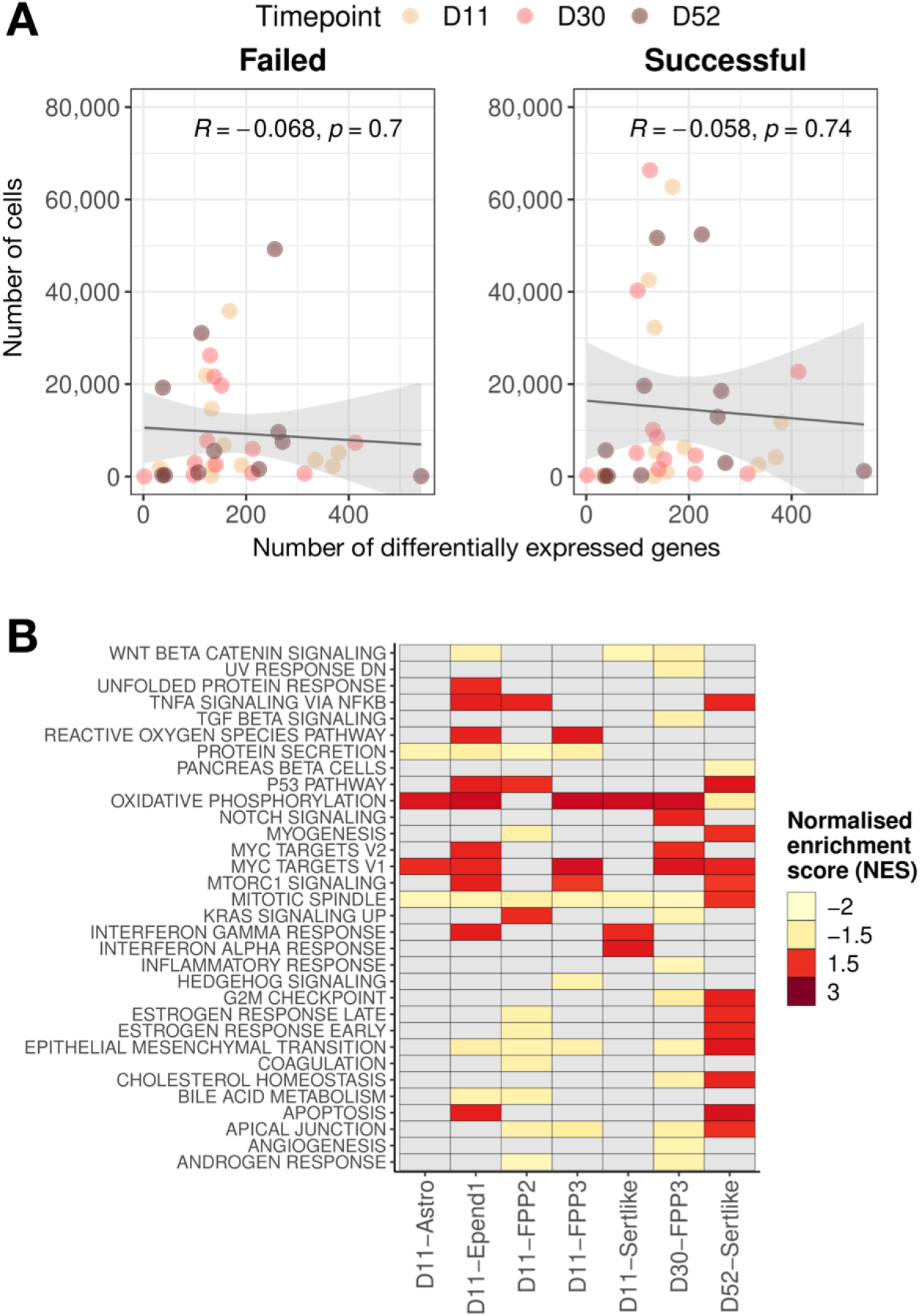
**a**, The number of cells per outcome and cell type did not correlate with the detected number of differentially expressed (DE) genes (p>0.05), suggesting that differential gene expression was not biased by the sample sizes. Each dot represents a cell type and is coloured according to the corresponding time point. **b**, Gene set enrichment analysis on MSigDB hallmark signatures (|NES|>1.5, adj.P<0.05) among those cell types with cancer-associated differential gene expression. Pathways with significant enrichment are coloured in red for upregulation, in yellow for downregulation and in grey when no significant enrichment is observed. Astrocytes at day 52 downregulated the oxidative phosphorylation and upregulated the mitotic spindle assembly, contrary to other cell types.

**Supp. Fig 5.**
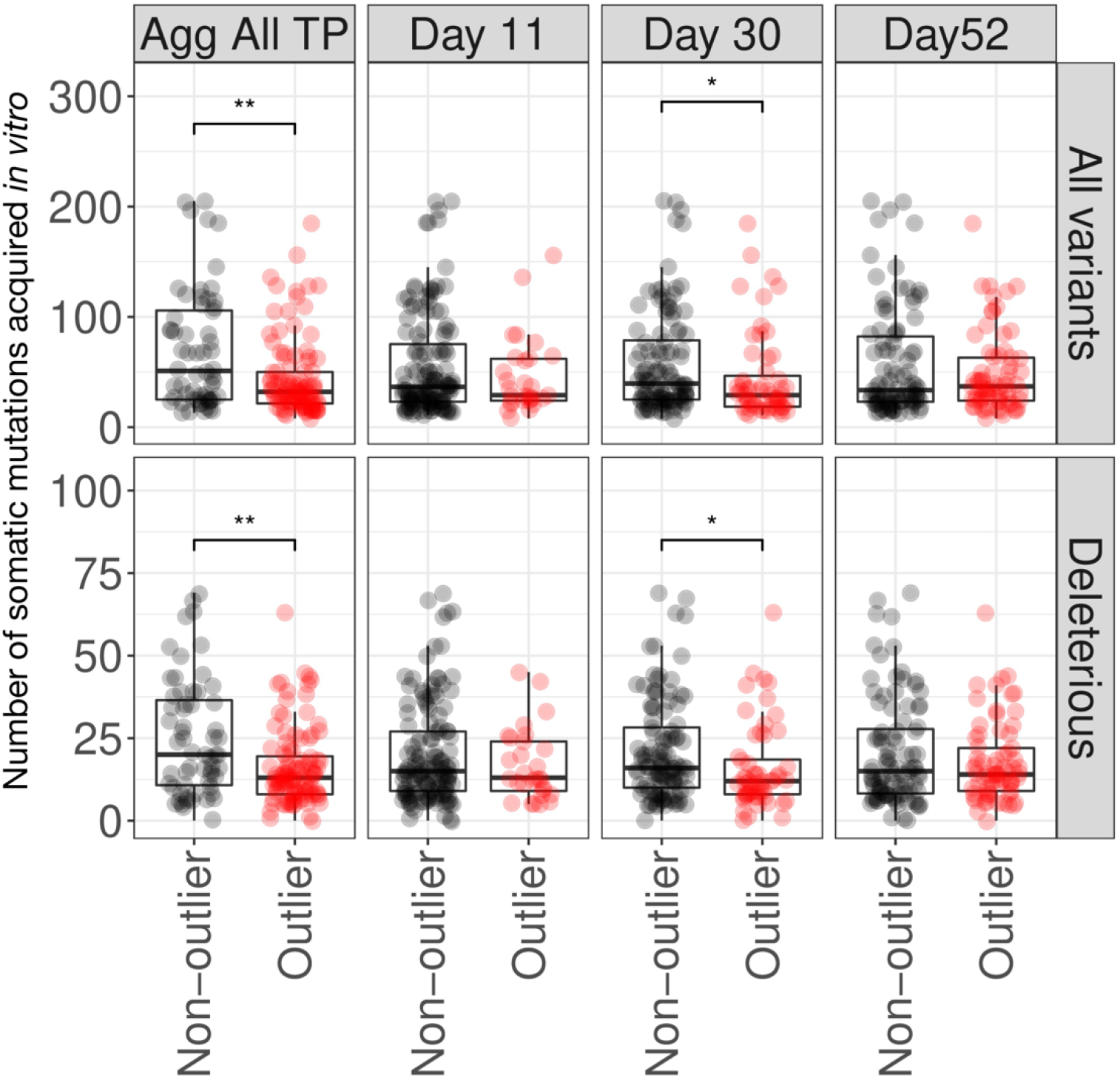
The differences in the burden of somatic mutations acquired *in vitro* between outliers and non-outliers of cell type composition are not consistent throughout the different stages of the differentiation (Wilcoxon Rank Sum Test). The definition of outlier lines was based on the observation of an abnormal cell type proportion during the differentiation or at a given time point. Only at day 30, we observed a reduced burden in the outlier group, which was also observed with the aggregated definition, both when accounting for total variants or for deleterious only.

## Materials and Methods

### 1. Quality control and annotation of somatic mutations acquired *in vitro*

A joint variant calling (BCFtools/mpileup and BCFtools/call, version 1.4.25, human genome assembly GRCh37d5) between 384 pairs of parental fibroblasts and their corresponding iPSC lines was performed to identify 18,999 somatic mutations that were acquired or positively selected throughout iPSC reprogramming as described in ^11^. This calling was performed for single-nucleotide variants (SNVs), dinucleotides, indels and copy number variants (CNVs) for both autosomal chromosomes and chromosome X. Only in the case of indels, chromosome X was not included.

We filtered the initial call set ^11^ following these steps: the variants out of the exome sequencing baits were excluded, germline variants were filtered-out assuming they show a minor allele frequency MAF>0.1% in 1000 Genomes Phase3 ^40^ or in ExAC 0.3.1 ^41^, or be carried by the parental fibroblast of more than one donor. Only high-quality variants were filtered-in (PASS filter). Variants with an allelic fraction larger than 0.6 in iPSC or in fibroblasts were removed to filter potential spurious mutation calls. We classified variants either as acquired *in vitro* or positively selected only when a significant rise in allele frequency was observed between the iPSC and the parental fibroblast (Fisher’s exact test p<1.6·10^−4^, equivalent to FDR 5% using the Benjamini-Hochberg multiple test correction procedure). 17 out of the 384 iPSC lines were found to be hypermutated *in vitro* (>240 mutations corresponding to a Z-score>2) and were discarded for further analysis downstream. Finally, we defined our gene universe as the 19,653 genes that were protein-coding genes in the Ensembl gene annotation (GRCh37, version 87) and were covered by the exome sequencing baits. In line with that, we discarded all those variants that could not be annotated to the gene universe, finally releasing a call set of 18,999 mutations, including 460 CNVs, 642 indels, 2,445 dinucleotides and 15,452 SNVs (**Supp. Table 1**).

### 2. Functional annotation of somatic mutations acquired *in vivo*

We annotated the somatic mutations acquired *in vivo* (SNVs, dinucleotides, indels) by predicting their functional consequence using the variant effect predictor (VEP, release 99) ^42^ and the haplotype-aware BCFtools/csq tool (version 1.9) ^43^. We used Ensembl gene annotations (GRCh37, version 87) and recorded only the most impactful consequence for each mutation, as determined by the following decreasing order of severity: https://www.ensembl.org/info/genome/variation/prediction/predicted_data.html

We defined mutations as loss-of-function (LoF) when they were annotated as frameshift, stop-gain, splice acceptor or splice donor variants; and as missense pathogenic (or damaging missense) when annotated as missense or start loss with a CADD Phred score cutoff > 15 (version 1.6) ^44^. The definition of deleterious mutations included the union of LoF and missense pathogenic mutations. The remaining mutations were annotated as synonymous or as “others” (either as coding, non-coding or unannotated). Overall, we annotated 1,002 LoF mutations, 5,722 missense pathogenic, 2,631 missense non-pathogenic, 3,158 synonymous and 6,026 other mutations (**Supp. Table 1**).

### 3. Definition of differentiation outcomes in iPSC-derived cell types

#### Sensory neurons ^14^

We processed the Supplementary Table 1 of the original publication (“IPSDSNs”) that contained the metadata for all cell lines differentiated to sensory neurons. We annotated the differentiation outcome by re-labelling the neuron quality status per line from “Poor” to failed and from “Good” to successful. We excluded those cell lines with undefined neuron quality status and renamed the non-neuronal differentiation outcome to failed differentiation. We identified 13 iPSC lines with differentiation replicates, all of which had a concordant outcome (2 failed, 10 successful), except for one line (“HPSI0613i-eojr_3”) that was discarded. In total, we annotated the outcome for 105 cell lines (5 failed and 100 successful), of which 85 lines (3 failed, 82 successful) were also profiled by WES and had mutation data available (**Supp. Table 1**).

#### Macrophages ^13^

We processed the Supplementary Table 1 of the original publication that contained the status of the differentiation outcome per line. We removed those cell lines that showed a low macrophage purity (“FC_QC_fail”) or presented degraded RNA (“RNA_QC_fail”). We identified 11 iPSC lines with differentiation replicates, all of which had a concordant outcome (7 failed, 4 successful). In total, we annotated the outcome for 123 cell lines (23 failed and 90 successful), of which 102 lines (22 failed, 80 successful) were also profiled by WES (**Supp. Table 1**).

#### Dopaminergic neurons ^8^

We reprocessed the whole scRNA-seq dataset (excluding cells treated with rotenone), re-clustering all cells at once and annotating the resulting cell types using the same markers from the original publication (see also **Section 7**). We then computed the total number of cells and the cell type proportion per line in each pool. For each time point, we removed those cell lines with the lowest number of cells (first twentile) to increase the confidence level of the cell type proportion estimates. We then defined the neuronal differentiation efficiency per cell line in each pool as the proportion of dopaminergic and serotonergic neurons observed at day 52 of the differentiation. We annotated this efficiency for 209 cell lines distributed in 18 pools, and classified lines accordingly either as failed lines (<0.2, this includes lines with poor/impaired outcome) or successful lines (≥0.2). Only 3 out of the 36 iPSC lines placed in more than one pool (pool replicates) were discordant. The DN outcomes for the remaining 206 lines corresponded to 56 failed and 150 successful lines, with a subset of 126 lines also profiled by WES (35 failed, 91 successful). Overall, we reached a 98.8% agreement with the neuronal differentiation outcome classification of the original paper.

Additionally, we processed the Supplementary Table 5 ^8^ which contained the model scores from the predicted efficiency. Those scores were obtained from a logistic regression trained with a binary outcome definition per line (either successful lines with >20% measured efficiency or failed lines with <20%) and an independent dataset of bulk RNA-seq that uses all expressed genes from 184 iPSC lines. The model scores classified 812 HipSci iPSC lines as failed (N=103) and successful (N=709) differentiators (precision=0.9 and recall=0.35 for threshold=0.02231), with 349 lines also profiled by WES (33 failed, 316 successful) (see **Figure 1A, Supp. Table 1**).

#### Endoderm ^15^

We processed a table obtained from the authors of the original publication with the differentiation efficiency for 108 donors, of which 86 were also profiled by WES. Here, differentiation efficiency is computed as the average pseudotime on day 3, having a continuous distribution of efficiencies rather than a binary outcome (“failed”, “successful”).

### 4. Gene burden differences linked to the differentiation outcome

We leveraged the 832 iPSC lines profiled with WES available from the HipSci project and annotated the most severe consequence for each variant using the variant effect predictor (VEP, release 99) ^42^ and the Ensembl gene annotation from the release (version 75). We summarise the results by building a matrix of mutation counts per variant category either for LoF or deleterious mutations representing gene damage categories, or synonymous variants as mutational burden control. The criteria for inclusion in each of the categories was the same as provided previously with the annotation of somatic acquired mutations *in vivo* (see **Section 2**). Each matrix contained the 19,653 genes (of the gene universe) as rows and the 832 lines as columns.

We then combined the mutational burden data per line for each variant category (LoF, deleterious and synonymous) with the corresponding binary differentiation outcome (DN predicted (N=793), DN observed (N=183), macrophages (N=118)). We excluded the sensory neurons from the analysis due to the low number of failed lines in that dataset (N=3). For each variant category and outcome combination, we performed a Wilcoxon Rank Sum Test per gene (N=19,653 tests) to identify those that presented a differential burden between failed and successful lines. For each combination, we performed a multiple test correction using the Benjamini & Hochberg approach. The level of statistical significance was set at FDR=5%. To compute a fold change of the mutational differences per gene between failed and successful lines, we initially normalised the mutation ratio by the gene length (as in Ensembl Annotation release 87) and the number of lines per outcome and divided the ratios using pseudocounts. The pseudocounts used for each combination corresponded to the minimum non-zero normalised mutation rate observed across all genes for any of the outcomes. We used a threshold of FC>2.5 or FC<1/2.5 to classify genes as disproportionately mutated in failed lines or in successful lines, respectively. Alternatively in the case of the DN dataset, we also combined the burden data with the continuous distribution of differentiation efficiencies and predicted scores to check the robustness of the *BCOR* association. In this case, we performed a Pearson’s correlation per gene for each combination.

### 5. Gene ontology enrichment analysis for DN differentiation

For the gene ontology enrichment analysis on biological processes, we only focused on the deleterious burden of iPSC lines. Given the number of lines for the DN differentiation (183 for the observed outcome and 793 for the predicted one), it is the variant category linked to gene damage with the best power to detect enrichment. We proceeded with the matrix of counts previously generated for the identification of gene burden differences linked to the differentiation outcome. Likewise, we computed the fold change of mutational differences per gene between failed and successful lines using the ratio of the mutation burden (mutations per Kb) normalised by gene length, number of lines per outcome and adding 0.001 as pseudocounts. Prior to the GO analysis, we annotated all genes with their corresponding Entrez gene identifiers and selected the most differentially mutated genes in successful lines (top-5% in log_2_FC) and the most differentially mutated in failed lines (bottom-5% in log_2_FC). We then run the hypergeometric test for GO term overrepresentation of biological processes conditional to the hierarchical GO structure (package *GOstats* from R). For each of the four tests, we provided the selected genes in each DN outcome combination: failed/observed, failed/predicted, successful/observed and successful/predicted. We used the following thresholds: the cutoff for significance was set at p<0.05, we considered only those gene sets defined with more than 20 genes and each gene set had to account for at least 10 counts in each analysis. All gene sets found to be significantly enriched are shown in **Supp. Table 2**. Finally, we highlighted only those significant GO terms related to neurodevelopment or chromatin modification, so we highlighted any gene set with the following words in **Fig. 2D:** *“Axon”, “neuron”, “glial”, “brain”, “hindbrain”, “forebrain”, “midbrain”, “synapse”, “chromatin”, “cerebellum”, “neural”, “cortex”, “neurogenesis”, “axonogenesis”, “nervous”, “hippocampus”, “neurotransmitter”, “dopaminergic”, “axenome”, “action potential” and “synaptic”*.

### 6. Gene set enrichment (DDD and Cosmic-Tier 1) in differentially mutated genes from discordant replicate lines

We identified 49 replicate line pairs with a discordant DN outcome, that is when one of the lines from a donor was predicted to fail differentiation (model score < 0.02231) and the other was predicted to produce dopaminergic neurons successfully (model score > 0.02231). One of the 49 pairs was added from a quartet of replicate lines (“HPSI-fdpl” donor), from which we selected the failed line and randomly sampled one of the three successful lines.

For each of the replicate pairs, we filtered out variants that were shared by the two lines with the purpose to retain only the mutational burden specific to each outcome. With the remaining variants, we annotated the most severe consequence predicted by VEP (version 99), as described earlier. For further analysis downstream, we only considered those variants annotated as LoF, deleterious or synonymous variants, that represent gene damaging categories and a control group, respectively. We summarised the mutation counts per gene and variant category for the 49 replicates and computed the fold-change mutational differences between failed and successful lines (log_2_FC, pseudocounts=1).

We focused on two curated gene sets of key relevance in development: developmental disorders genes (from the Deciphering Developmental Disorders Project, DDD; version 2.2 from DDG2P) ^29^ and cancer-associated genes (Cosmic: catalogue of somatic mutations in cancer, version 90 on GRCh37). In both sets, we filtered out those genes not overlapping our gene universe and considered only the DDD genes with monoallelic requirement (N=1,938), as well the Cosmic genes under the strongest oncogenic activity, labelled as Tier 1 (N=558).

We assessed whether the genes mostly mutated in failed lines (FC>2) for a given variant category are enriched in one of those gene sets when compared with those genes mostly mutated in successful lines (FC<½). We ran a proportion test with the mutational burden of that genes (2×2 table, rows: failed/successful lines, columns: gene set / non-gene set) and ranked the resulting p-value with those obtained from 1,000 random gene sets to finally compute an empirical p-value (i.e., position N would correspond to p=N/1000). Significant enrichment was considered only when empirical p<0.05.

### 7. Reanalysis of the pooled single-cell data of dopaminergic neuron differentiation Sample

#### selection and data preprocessing

The dopaminergic neuron differentiation was profiled by droplet-based scRNA-seq (10x Genomics). We processed a subset of the dopaminergic neuron differentiation dataset ^8^ that consisted of 119 10x samples out of the total 166 (**Supp. Table 4**). Here, a 10x sample is defined as the cells sequenced from one inlet of a 10x chip. For the sake of this study, we did not include those samples from the original experiment profiled under rotenone treatment at day 52 or containing iPSC-derived cerebral organoids (day 119). We also did not process samples for pool 10 (day 11) due to reported problems on library preparation.

We processed the 119 10x samples using CellRanger software (version 3.1.0) and aligned them to the GRCh37/hg19 reference genome. Gene counts were quantified by the “count” option of the software, using the Ensembl 87 reference gene annotation (N=32,738 genes). After pre-processing, we excluded 4 additional 10x samples due to quality control issues, mainly due to low percentages of cell singletons in deconvolution (≤50%) and low cell viability: two technical replicates from pool 12 on day 52, one sample from pool 8 on day 30 and a sample from pool 1 on day 30 (**Supp. Figure 2b**). The final 115 10x samples covered all pooled experiments (N=19, pools 1-17 and pools 20-21), including 238 different cell lines (7-24 lines per pool). Only one cell line (“HPSI0913i-gedo_33”) was removed due to an abnormally high cell line proportion (>90%) in pool 14.

#### Quality control and deconvolution of cell donor identity

Each 10x sample went through a quality control step in which we removed dying cells or those with broken membranes, displaying a low number of genes per cell (<200) and an excess of mitochondrial count fraction (>5%). Also, we discarded those cells with an abnormal percentage of reads consumed by the top-100 mostly expressed genes (<75%), which indicate technical artefacts affecting the good coverage of the full transcriptome of the cells. On the other hand, we filtered out those genes that were not expressed in at least 0.1% of the total cells.

For each of the 19 pools, we performed cell deconvolution using *demuxlet* ^45^ using existing genetic variation (genotypes of common biallelic exonic variants, MAF>5%) available from the HipSci Project ^7^ as in the original paper. In those cases when iPSC cell lines had not been genotyped (intended cell lines from **Supp. Table 3**), we used the genotypes from the primary fibroblast instead when available. Demuxlet was run using a default prior doublet rate of 0.5. We only retained those (singletons) cells that could unambiguously be linked to a donor and discarded those 10x samples for which the overall singleton outcome was low (<50%) (**Supp. Figure 2b**). We performed the quality control and the integration of the cells from the 10x samples using the Scanpy Python-based toolkit (version 1.4.5.1) ^46^.

#### Normalisation, dimensionality reduction, batch correction and clustering

We performed a combined analysis of all the three time points (day 11, day 30 and day 52) to have a shared embedding for all lines. Initially, genes that were not expressed in at least 0.1% of total cells were removed. Then, gene counts were normalised to the total number of counts per cell and log-transformed (log1p). After adjusting for mean-variance dependence, we selected the 2,928 highly variable genes and scaled gene counts to unit variance and zero mean. We then calculated the first 50 principal components (PCs) and batch-corrected them with Harmony ^31^, treating each 10x sample as a different batch (parameters: theta=2, max.iter.harmony=25, max.iter.cluster=500). We then used the batch-transformed PCs to compute a neighbourhood graph (n_neighbors=10), visualise it using UMAP and perform the cell clustering using the Leiden algorithm (resolution=0.3) identifying 12 different clusters (**Supp. Figure 2h-j**). We also used the Scanpy toolkit (version 1.4.5.1) for all the steps, except for Harmony that was run in R version 4.0.3.

#### Cell type annotation

Cell type annotation was performed using the same set of literature-curated markers as in ^8^ (**Supp. Figure 2e-g**). We confidently annotated 10 out of the 12 identified clusters. Interestingly, we could also identify neuroblasts at day 11 with a commitment to dopaminergic neurons, as they clustered together, but at the same time did not exhibit the neuron marker expression.

At day 11, we characterised four populations of floor-plate midbrain progenitors, either proliferating (proFPP-1, proFPP-2) or non-proliferating (FPP-1, FPP-3). We could also link the population of neuroblasts to their early dopaminergic or serotonergic commitment. At day 30 and 52, we identified six additional cell types, including another cell type of non-proliferating progenitors (FPP-2), two neuronal populations (dopaminergic-like (DA) and serotonergic-like (Sert-like) neurons) and three non-neuronal ones (astrocytes (Astro), ependymal-like cells (Epend-1) and an unknown population (Unk-1) potentially linked to Cajal-Retzius transient neurons. At day 30, two additional rare cell types (<2%) were identified, either belonging to a subgroup of Sert-like neurons associated with proliferation markers (pro.Sert-like), or to an unknown population (Unk-2) only detected in a single 10x sample of pool 12.

### 8. Reproducibility of cell line abundance in different types of replicates

We started processing the metadata object containing annotations from all cells (**Supp. Data 1**). To define different types of replicates, we used the information provided by the naming of the 115 10x samples (**Supp. Table 4**). Four replicate types were defined per cell line (**Supp. Fig. 3a**):

**- Pool replicates**: The same cell line was placed in different pooled experiments (different cell lines in the background).

**- Biological replicates**: One cell line underwent independent differentiations (different time and plate), but within the same pool (same background).

**- Technical replicates**: One cell line underwent differentiation within the same pool, same time, but different wells of the same plate.

**- 10x replicates**: One cell line underwent differentiation within the same pool, same time and same well of the plate.

Initially, we calculated the cell line proportion within each 10x sample. Then, for each cell line, the corresponding replicate group and a given time-point, we calculated the averaged cell line proportion per replicate taking into account the contributing 10x samples. For instance, to compare the cell line proportion of the two biological replicates for “HPSI1014i-tuju_1” at day 11, we averaged the cell line proportion of the four 10x samples contributing to replicate 1 on one side, and the four 10x samples contributing to replicate 2, on the other. To evaluate the reproducibility between replicate proportions, we fitted a linear regression and computed the adjusted R-squared and the p-value of the association. Data points on **Fig. 3A** corresponds to matched replicates per line/time-point combination. Note here that the designation of replicate 1 or replicate 2 before the regression is random.

### 9. Cell line proliferation throughout dopaminergic neuron differentiation

The cell suspension for each pool was prepared with an equal amount of each iPSC line ^8^. For this analysis, we only considered those cell lines within a given pool that had been sampled in the three time points of the differentiation (N=187). Given their lack of reproducibility, pool replicates (N=23) were considered as independent lines. For each pool and time-point, we computed the log-transformed (log1p) cell line proportion. Then, we calculated the proliferation rate at day 11, day 30 and day 52, dividing the cell line proportions observed at each respective time point by the equal proportions from day 0. We then annotated each cell line with the observed outcome in DN dataset, either successful or failed, based on the neuron differentiation efficiency threshold of 0.2. Additionally, we annotated failed cell lines with either *BCOR+* (N=23) or *BCOR-* (N=33) based on the presence of LOF *BCOR* variants in each line. Note here that none of the successful lines (N=131) carried any LoF mutation in our iPSC exome-sequencing data.

### 10. Annotation of cancer-driver mutations

For each of the 832 iPSC cell lines with exome sequencing data, we annotated the most severe consequence for each variant using the variant effect predictor (VEP, release 99) as described earlier (see **Section 2**). We then overlapped the predicted LoF variants from each line with those listed in the database for the cancer-associated genes (Cosmic Tier 1, version 94) under the strongest evidence for oncogenic activity (Tier 1). We restricted the overlap search to those Cosmic Tier 1 variants that have a defined genomic position and a FATHMM score >= 0.7 ^11^. We identified 726 potential driver mutations with 606 iPSC lines carrying at least one driver mutation. The most mutated driver genes were *PDE4DIP* (564), *CCND3* (35), *TCF3* (29) and the *BCOR* (24), also after normalising the burden by the CDS length. We then annotated each cell line with both the observed and predicted DN differentiation and computed the driver mutational ratio (log2-transformed) given the outcome. In both cases, *BCOR* ranked as the gene with the highest ratio of driver mutations in failed lines versus successful ones (5:0 with the observed DN outcome, 18:5 with the predicted DN outcome).

### 11. Cell type composition analysis between failed and successful lines

After annotating the cell type, we used the metadata information for each cell (**Supp. Data 1**) to compute the cell type proportions per line. We used the DN outcome annotation for the 206 lines classified either as failed or successful, as described in **Section 3**. For 8 pool replicates missing day 52 time-point in one of the pools, we imputed the same outcome as observed in the other replicate. We also included 15 out of the 18 lines that were not profiled at day 52, but with data from previous time-points (either from day 11 or 30, or from both). To take advantage of those lines in the cell type composition analysis, we classified them either as successful or failed using the predicted model scores from ^8^. Following those steps, we end up annotating the differentiation outcome for 221 lines (58 failed, 163 successful).

For each time-point and cell type, we used a negative binomial regression model to evaluate the composition changes between failed and successful lines. We modelled the total number of cells per line as an offset variable, given that the accuracy of cell type proportion estimates increases with their magnitude.

~~~
glm.nb(nCells ∼ outcome + offset(log(nTotalCells)))
~~~

We finally performed a multiple-test correction (N=36 tests) using the Benjamini & Hochberg approach (FDR<5%). The significance level of the mean cell type proportion difference between mutated and unmutated groups is indicated by different levels: pAdj<0.05 (*), pAdj<0.01 (**), pAdj<0.001 (***).

### 12. Association of cell type composition with *BCOR* mutational carrier status and in vitro proliferation rate

We calculated the cell type proportion estimates for the 209 lines on day 52 from the DN dataset (**Supp. Data 1**) as described in **Section 11**. For each gene (N=19,653), we compared the cell type composition differences at the end of the differentiation between those lines carrying at least one deleterious mutation (≥1) and those lines unmutated (Wilcoxon Rank Sum Test). For each cell type, we applied multiple-test correction (Benjamini & Hochberg, FDR<5%) to the raw p-values obtained from each gene and indicated whether they are more abundant in mutated (blue) or unmutated lines (orange). Note here that only those cell types that show >2% of abundance at day 52 are considered, discarding FPP-1, FPP-3, proliferative serotonergic-like neurons, proliferative FPP-2 and the unknown cell type 2.

Alternatively, for each major cell type at day 52 (>2% abundance), we tested for the association between cell type proportions per line and their corresponding proliferation rates between day 52 and day 0 (see **Section 9**) using Pearson’s correlation. We corrected for multiple-test correction using the Benjamini & Hochberg method (N=7, FDR<5%). We also indicated if the cell type proportions correlate (blue) or anti-correlate (orange).

The significance level of all comparisons was indicated as follows: pAdj<0.05 (*), pAdj<0.01 (**), pAdj<0.001 (***).

### 13. Differential gene expression analysis between failed and successful lines

We leveraged the gene expression data from 221 lines annotated with the DN outcome (58 failed, 163 successful) as in the cell type composition analysis. Overall, we processed 273,804, 266,226 and 306,811 cells from day 11, day 30 and day 52, respectively. For each time-point, we load the “.h5” object with the log-transformed gene expression per cell (log1p normalised counts, not-scaled) (**Supp. Data 2-4**) and filtered out those genes expressed in less than 1% of the cells, finally processing 12,912, 14,149 and 14,737 genes, respectively. We performed differential gene expression analysis between those lines that failed differentiation and those lines that were successful for each cell type and time point combination using the Wilcoxon Rank Sum test (as implemented in Seurat). We required more than 10 cells to be represented in each of the outcomes to run the DE test for each combination. DE genes were selected based on an adjusted P-value < 0.05 and a FC >1.5.

For each cell type and time point, we also tested the overrepresentation of three gene sets (Cosmic-Tier1, DDD and a subset of dominant DDD genes) among the list of DE genes (Chi-squared test using p-values computed by Monte Carlo simulation using 100,000 replicates). The gene universe for the test used the union of the pass-filtered genes across all timepoints (N=15,367). We corrected for multiple-test correction using the Benjamini & Hochberg method (N=102, FDR<5%). The significance level of each test was indicated as follows: pAdj<0.05 (*), pAdj<0.01 (**), pAdj<0.001 (***).

### 14. Gene ontology enrichment analysis of differentially expressed genes per outcome and cell type

We performed 12 gene ontology enrichment tests on biological processes based on:

- The union of genes found to be differentially expressed in any given time point and cell type combination (labelled as **allDE**).

- The union of genes found to be differentially expressed in any given time point and cell type combination with an overrepresentation of DDD DE genes (labelled as **allsignif**).

- The lists of DE genes for each of the 10 time point and cell type **combinations** with an overrepresentation of DDD DE genes (cyan circles with p<0.05, Fig. 4C).

Previous to the GO analysis, we annotated each feature of the gene universe (N=15,367) with their corresponding Entrez gene identifiers using the package *org*.*Hs*.*eg*.*db* from R/Bioconductor. We discarded from the analysis those genes without correspondence or showing duplicate identifiers. We then run the hypergeometric test for GO term overrepresentation of biological processes conditional to the hierarchical GO structure (package *GOstats* from R). Given the different magnitude of DE genes between **allDE** (N=1,884) or **allsignif** (N=972) and each of the combination tests (N=131-372), we used different thresholds for significance in each case: **allDE/allsignif:** {minimum gene set size = 30, maximum gene set size=200, pAdj<0.001, minimum number of counts per gene set = 20}; **combinations:** {minimum gene set size = 10, maximum gene set size=200, pAdj<0.001, minimum number of counts per gene set = 7}

All gene sets found to be significantly enriched are shown in **Supp. Table 5**. Finally, we highlighted only those significant GO terms related to neurodevelopment or chromatin modification, so we highlighted any gene set with the following words in **Fig. 4C:** *“Axon”, “neuron”, “glial”, “brain”, “hindbrain”, “forebrain”, “midbrain”, “synapse”, “chromatin”, “cerebellum”, “neural”, “cortex”, “neurogenesis”, “axonogenesis”, “nervous”, “hippocampus”, “neurotransmitter”, “dopaminergic”, “axenome”, “action potential” and “synaptic”*.

### 15. Gene set enrichment analysis on cell types with differential expression of cancer genes

We performed a gene set enrichment analysis (GSEA) on those time point and cell type combinations in which the list of differentially expressed genes were enriched in cancer-associated genes. For this purpose, we used the curated MSigDB hallmark gene set signatures (version 7.4 for symbol identifiers) ^47^. We considered only those gene sets with a larger size than 10 genes. For each gene set, we then ran the preranked gene set enrichment analysis with a maximum gene set size of 500 genes, an eps parameter of 0 and used 10,000 permutations for preliminary estimation of p-values. We highlighted significantly enriched gene sets as those with a BH-adjusted p-value < 0.05 and an enrichment score normalised to mean enrichment of random samples of the same size (NES): NES≥ 1.5 for upregulated pathways and NES ≤ -1.5 for downregulated ones.

### 16. Identification of cell line outliers for cell type composition

We calculated the cell type fraction per line within each pool and time point, removing those lines with the lowest number of cells (first twentile). Those cell lines pooled in more than one experiment (pool replicates) were considered as independent lines. We then computed the z-score associated with the calculated fractions and marked as outliers those lines showing a cell type fraction with a |Z-score|>2, either showing a deficiency or an excess of a given cell type.

We characterised the outlier behaviour focusing on those outlier cell lines represented in the three time points of the differentiation (N=121). Based on that, we computed the number of times each line shows an abnormal cell type fraction (outlier event) throughout the entire differentiation or specifically per time point. For each cell line, we explored when outlier events occurred and identified the most common time point combinations.

For those lines profiled with WES for the iPSC and the corresponding parental fibroblasts, we annotated the burden of somatic acquired mutations *in vivo* (**Supp. Table 1**) considering either total or deleterious variants. We then evaluated whether cell lines defined as outliers of cell type composition showed mean differences on the mutational burden (Wilcoxon Rank Sum Test). Alternatively, we also evaluated the outlier lines within each specific time point (day 11, N=58; day 30, N=82; day 52, N=114).

Finally, we also annotated each pooled cell line with their corresponding proliferation rate at day 52 (see also **Section 9**) and fitted a logistic regression to predict the outliereness of cell type composition.

### 17. Association between gene expression and cell type abundance

We analysed the correlation between the cell type abundance and the cell type specific expression for all genes using the existing cell line variability throughout the differentiation. We computed the z-scores for cell type composition as shown in **Section 16**. As for the gene expression (log1p normalised counts, not-scaled), we computed the z-score per cell type using the average gene expression per line at each time point. The average gene expression per line was calculated considering all the cells of that given line in one specific pool experiment, including those from different 10x samples when available. We required a minimum of 10 cells per line (in any time point - cell type combination) to calculate the average gene expression. We discarded all those combinations in which less than 10 lines matched this threshold (proliferative Sert-like neurons at day 11 and the glial cells (Unk-2) at days 11 and 52).

We then correlated the expression z-scores with the cell type fraction z-scores, as shown in the example for *KMT2D* gene for DA and FPP-1 in day 11 (**Fig. 5b**). To identify the key genes driving the outliereness in cell type composition, we performed the z-score correlations for all the detected genes per time point (day 11, N=12,912; day30, N=14,149; day 52, N=14,737). Those genes with no detectable expression in at least 10 lines of a given combination were not considered. We then sampled the genes with either positive or negative significant associations (p.Adj<0.05) from the resulting distribution of Pearson correlation coefficients per cell type (**Fig. 5c**, lower).

From the list of genes with significant correlation (or anti-correlation) per cell type and time point, we tested whether there was a gene set enrichment on key developmental genes, for both DDD and Cosmic-Tier1 genes. In this case, we performed a 2×2 chi-squared contingency table between the list of significantly and non-significantly correlated genes and the overlap with each gene set separately (Monte Carlo simulation using 2,000 replicates to compute the p-values). We then performed multiple-test correction using the Benjamini & Hochberg method (N=66 tests, FDR<5%). The significance level of each test was indicated as follows: pAdj<0.05 (*), pAdj<0.01 (**).

## Data availability

### Exome sequencing data from the HipSci Project

Links to the raw data are available from the HipSci project website (www.hipsci.org) for both the parental fibroblasts and the iPSC lines. Exome sequencing data for the open access samples is deposited in the European Nucleotide Archive (ENA, https://www.ebi.ac.uk/ena/) under the ERP006946 study accession number (N=325 iPSC lines, of which 260 also available for the parental fibroblasts). WES data for the managed access samples is deposited in the European Genotype-Phenotype Archive (EGA, https://ega-archive.org), with normal and specific disease cohort datasets available upon request and data access agreement. The variant call sets of acquired mutations *in vitro* and the code used to generate them are available at ^11^.

### Single-cell profiling of iPSC dopaminergic differentiation

Managed access data from scRNA-seq are accessible in the European Genome–phenome Archive (EGA, https://www.dev.ebi.ac.uk/ega/) under the study number EGAS00001002885 (dataset EGAD00001006157). Open access scRNA-seq data are available in the European Nucleotide Archive (ENA, https://www.ebi.ac.uk/ena/) under the study ERP121676(https://www.ebi.ac.uk/ena/browser/view/PRJEB38269). Processed single-cell count data and metadata tables (**Supp. Data 1-4**) will be made available on Zenodo. Chip genotypes for HipSci lines are available from the EGA (EGAS00001000866) and the NCBI (PRJEB11750).

## Code availability

All scripts and figures will be made available in the following github repository: https://github.com/paupuigdevall/somaticBurdenNeuro2022. The repository includes the single-cell processing scripts, the downstream analysis, and an R markdown with the main figures, as well the supplementary ones.

## Supplementary Information

### Supplementary Data

**Supplementary Data 1:** Metadata information for the 846,841 processed cells from the DN dataset: donor identity, cell type annotation, pool identifier, 10x sample, time point and replicate information.

**Supplementary Data 2-4:** AnnData/H5AD files containing the single-cell gene expression matrices and the metadata for day 11, day 30 and day 52, respectively. The gene expression is normalised and log-transformed, but not scaled.

**Supplementary Tables**

Supplementary tables are available at the downloadable file

`suppTableMutBudenNeuro.xlsx`.

**Supplementary Table 1**. Cell lines annotated with the burden of somatic mutations acquired in-vitro and their outcome in multiple differentiations (sensory neurons, macrophages, dopaminergic neurons -DN- and endoderm). 842 HipSci iPSC cell lines are included.

**Supplementary Table 2**. Results from the gene ontology enrichment analysis for the most differentially mutated genes (top-5% in deleterious mutations) per dopaminergic differentiation outcome. Both predicted (N=793) and observed (N=183) DN outcomes are included.

**Supplementary Table 3**. List of the cell lines used in each of the 19 pool experiments from the DN dataset. **Supplementary Table 4**. List of the 119 10x samples from DN dataset that were pre-processed with CellRanger. 115 out of the 119 samples passed the QC filtering criteria and were processed further downstream.

**Supplementary Table 5**. Functional enrichment analysis for the differential gene expression between failed and successful lines in the DN outcome. Results from the several hypergeometric tests for GO term association.

## Acknowledgements

This work was funded by the UK Medical Research Council (MR/L016311/1; MRC eMedLab Medical Bioinformatics career development award to HK), Helsinki Institute of Life Science (HK), the ICH CIO (SC and PP) and the NIHR GOSH BRC (SC, PP and HK). The views expressed are those of the author(s) and not necessarily those of the NHS, the NIHR or the Department of Health. The authors gratefully acknowledge Drs Daniel Gaffney, Oliver Stegle, and Florian Merkle for early access to the dopaminergic neuron single-cell dataset, and Drs Daniel Seaton and Anna Cuomo for assistance with the DN data. The authors also acknowledge Drs Matthew Hurles, Sebastian Gerety and Osama Arshad for their feedback on the project.

## Notes

### Competing Interest Statement

The authors have declared no competing interest.

